# Environmental controls on crenarchaeol distributions in hydrothermal springs

**DOI:** 10.1101/2024.07.09.602736

**Authors:** Amanda N. Calhoun, Jerome Blewett, Daniel R. Colman, Maximiliano J. Amenabar, Carolynn M. Harris, Eric S. Boyd, Ann Pearson, William D. Leavitt

## Abstract

Thermophilic archaea synthesize cellular membranes composed primarily of isoprenoid glycerol dibiphytanyl glycerol tetraethers (iGDGTs). Cells can adjust the packing of their lipids by increasing the number of cyclopentyl rings during lipid synthesis, thereby decreasing membrane permeability and fluidity to maintain cellular function at high temperature, acidic pH, or nutrient limitation. Archaea of the class *Nitrososphaeria* synthesize an iGDGT, crenarchaeol, with four cyclopentyl rings and a cyclohexyl ring, the function of which is unknown. Structural modeling suggests the cyclohexyl ring may increase membrane fluidity, potentially optimizing membranes for mesophilic conditions. To begin to investigate this hypothesis, iGDGT composition was quantified in forty-one thermal springs in Yellowstone National Park (YNP), USA, and contextualized within a global thermal spring iGDGT compilation with pH values of 1.1 to 10.1 and temperatures of 16 to 95°C. pH was the strongest predictor of both crenarchaeol relative abundance and the number of cyclopentyl rings per iGDGT. Crenarchaeol relative abundance exhibited a nonlinear relationship with pH and temperature, with highest relative abundances at pH 7.4 and 46°C, then decreasing above and below these values. These observations are consistent with the hypothesis that the cyclohexyl ring of crenarchaeol optimizes archaeal cellular membranes for circumneutral and moderate temperature environmental conditions.

**Importance:** Archaea change the composition of their membrane lipids to alter the fluidity of their membranes to maintain cell homeostasis when confronted with environmental stress. Some archaea of the class *Nitrososphaeria* produce a lipid, crenarchaeol, with a unique six-membered ring, the effect of which on archaeal membrane dynamics remains unknown. In this study, we identify pH as the most important geochemical variable for archaeal membrane response in Yellowstone National Park thermal springs. In addition, the lipid distributions indicate that crenarchaeol production is highest in circumneutral and mesophilic environments. The YNP results are supported by similar trends across global thermal springs.

## 1. Introduction

Terrestrial hydrothermal systems are valuable analogs of early Earth environments where many origin of life theories suggest life first emerged due to sharp thermal and geochemical gradients (Damer and Deamer, 2020). Archaea are ubiquitous in these environments and are adapted to the extreme conditions within these systems (van de Vossenburg et al., 1998; Macalady et al., 2004) that impose chronic energy stress (Valentine, 2007). A primary adaptation of archaea that inhabit extreme geothermal environments is the structural properties of their isoprenoid glycerol dibiphytanyl glycerol tetraether (iGDGT) lipids that comprise their cell membranes (van de Vossenberg et al., 1998; Konings et al., 2002). Archaeal iGDGTs are also ubiquitous in cells that inhabit soils, lakes, marine waters, and sediments, where they can serve as paleoenvironmental proxies (Schouten et al., 2000; Schouten et al., 2002; Schouten et al., 2013). Interactions between archaea and their environments have shaped their adaptive evolution (Colman et al., 2018; Yang et al., 2021), specifically the ability to adjust cell membrane fluidity and permeability in response to physical and chemical stressors (Oger and Cario, 2013).

The membranes of thermophilic archaea are primarily composed of iGDGTs (Macalady et al., 2004; Oger and Cario, 2013). These molecules vary in the number of cyclopentane or cyclohexane rings present in their internal biphytanyl chains (De Rosa and Gambacorta, 1980; De Rosa and Gambacorta, 1988; Schouten et al., 2000). The rings in iGDGTs affect molecular packing (Figure 1), enabling the modulation of archaeal cell membrane fluidity and permeability in response to changing temperature, pH, energy stress, salinity, and pressure through production of more or fewer rings as needed (Albers et al., 2000; Gabriel and Chong, 2000; Boyd et al., 2011; Pearson and Ingalls, 2013; Feyhl-Buska et al., 2016; Zhou et al., 2020;). In most environments, archaea produce iGDGTs with zero to four cyclopentane rings (iGDGT-0 to iGDGT-4), but in thermal springs, archaea can produce iGDGTs with up to eight cyclopentane rings (iGDGT-8) (Pancost et al., 2006; Schouten et al., 2013). The increase in average ring counts of iGDGTs at higher temperatures is the basis for the TEX_86_ (TetraEther indeX of 86 carbon atoms) paleotemperature proxy (Schouten et al., 2002), which is used to infer ancient environmental temperatures. However, recent studies indicate that additional environmental parameters can influence cyclopentyl ring abundance, complicating the interpretation of TEX_86_ as a strict temperature proxy (Qin et al., 2015; Elling et al., 2015; Hurley et al., 2016; Zhou et al., 2020; Cobban et al., 2020; Tourte et al., 2022).

**Figure 1.**
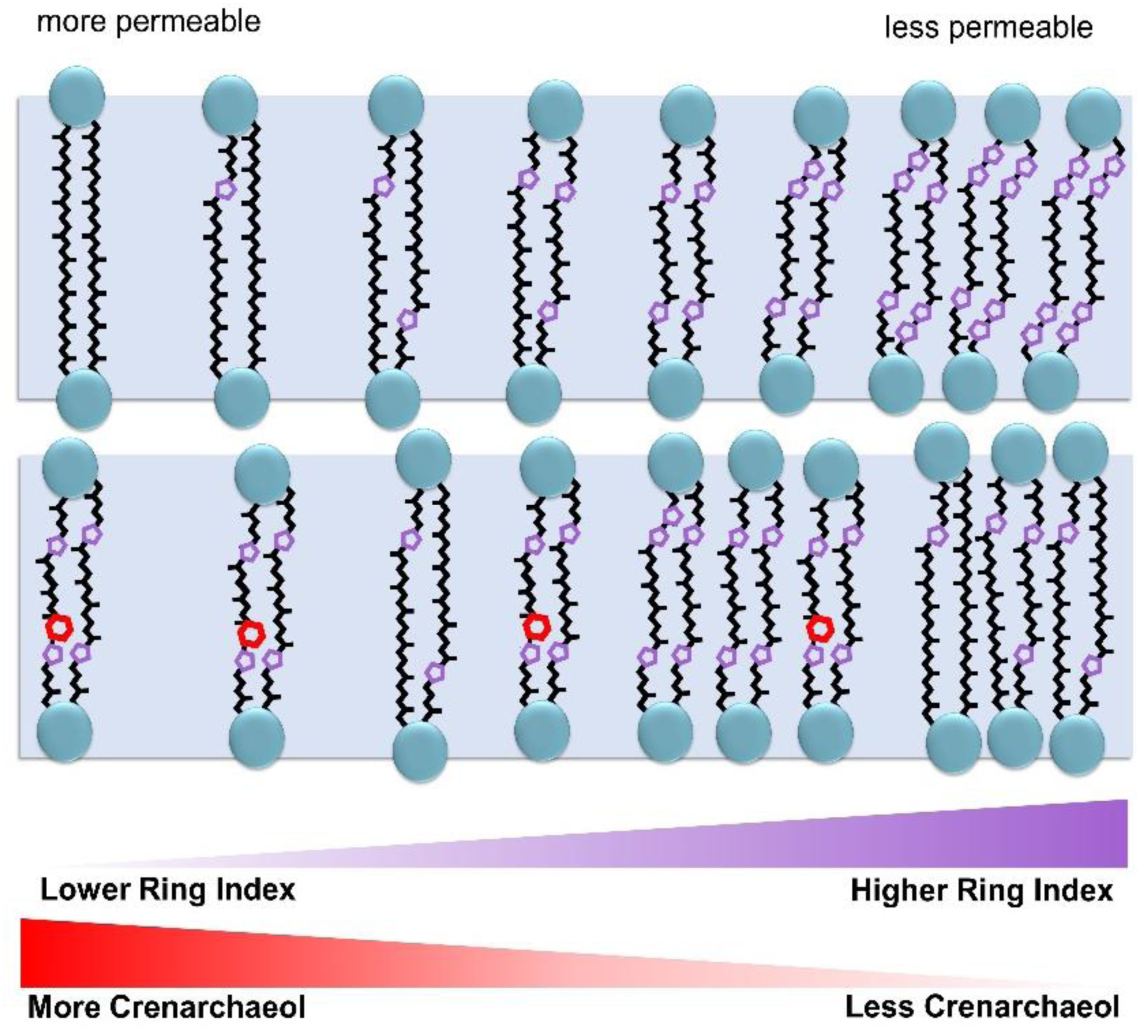
Schematic of effects of cyclopentyl rings (purple) on archaeal membrane permeability versus the hypothesized effect of cyclohexyl rings (red) on archaeal membrane permeability, indicated by lower bars.

Crenarchaeol iGDGT lipids and associated isomers are unique among iGDGTs in that they contain a cyclohexane ring (Sinninghe Damsté et al., 2002). Crenarchaeol is so far specific to representatives of the class *Nitrososphaeria* (syn. *Thaumarchaeota*) that are ubiquitous in terrestrial thermal springs, soils, and marine water columns where they can be the most abundant nitrifiers (Ueda et al., 1995; Karner et al., 2001; Stahl and de la Torre, 2012). At present, the origin and biosynthetic mechanism of this ring remain unknown, while hypotheses on its physiological function are not yet well-established (Zhang et al 2006). While recent work has revealed many steps of iGDGT synthesis (e.g., Zeng et al., 2022; Lloyd et al., 2022), the biosynthetic pathway of iGDGT cyclohexyl rings remains elusive, leaving the natural history and function of crenarchaeol unknown. In addition to crenarchaeol, organisms of the *Nitrososphaeria* produce iGDGT-0 to iGDGT-4, some of which contribute to TEX_86_ calculations. In thermal springs, the often-abundant *Nitrososphaeria* contain the only subgroup demonstrated to produce crenarchaeol, the ammonia-oxidizing archaea (AOA; De la Torre et al., 2008; Pitcher et al., 2010; Boyd et al., 2013). The crenarchaeol-producing AOA are thought to have originated as thermophilic descendants of non-AOA *Nitrososphaeria* (Abby et al., 2020; Luo et al., 2024). The predecessors of extant crenarchaeol-producing AOAs would have expanded their habitat from thermal springs to cooler marine waters between 629 and 412 Ma based on phylogenomic and marker gene (*amoA*) analysis of environmentally diverse archaeal genomes (Yang et al., 2021). As such, the progenitor that was originally optimized to live at high temperatures had to adapt its membrane structure away from the dense, rigid membranes of thermophilic archaea with high numbers of cyclopentyl rings. Incorporating the cyclohexyl ring into iGDGTs (i.e., crenarchaeol) may have relaxed the tight molecular packing in thermophile cellular membranes, thus facilitating the colonization of AOA progenitors in the less “extreme” marine realm (Schouten et al., 2000; Figure 1). This hypothesis is supported by lipid modeling that indicates the cyclohexane ring forms a bulge that increases biphytane volume, preventing dense membrane packing (Sinninghe-Damsté et al., 2002). A previous study suggested a 40°C temperature optimum for crenarchaeol given its abundance normalized to GDGT-0 in marine sediments and thermal spring microbial mats (Zhang et al., 2006), though crenarchaeol can be the most abundant core lipid in the membrane of cultured archaea above 70°C (de la Torre et al., 2008). While temperature is known to influence the membrane lipid composition of marine archaea, a clear negative correlation between crenarchaeol abundance and acidity in thermal spring archaeal communities (Boyd et al., 2013) implies that crenarchaeol is synthesized to a greater extent in circumneutral waters as opposed to acidic conditions. These competing predictions and observations motivate the current study to better understand whether temperature is the major determinant of environmental crenarchaeol distributions.

Here we utilize lipid data to examine patterns of crenarchaeol relative abundance and distribution from 299 thermal spring samples from North America, Europe, and Asia that span variable temperature (16 to 95°C), pH (1.1 to 10.1), redox (–330 to 330 mV for a new Yellowstone National Park 41 sample subset), and other geochemical conditions. In parallel, we calculate the summary indicator of cyclopentyl ring abundance, Ring Index (RI). This approach enables us to re-evaluate potential archaeal adaptations to environmental stress, allowing for the examination of key geochemical factors that drive crenarchaeol abundance on the present and past Earth.

## 2. Experimental Procedures

### Field Measurements and Sample Collection

A total of 41 sediment samples, encompassing 38 individual springs, were collected over four field seasons between 2018-2022 in Yellowstone National Park, USA (permit #YELL-05544) (Figure 2). Water temperature (°C), pH, dissolved oxygen (DO) concentration, specific conductivity (SPC), and oxidation reduction potential (ORP) were measured in the field at the time of sampling using a YSI ProDSS meter calibrated for DO and with pH buffer solutions of 4, 7, and 10 or 1.68, 4, and 7 (YSI, Yellow Springs, OH, USA). Concentrations of dissolved sulfide (ΣH_2_S/HS^-^/S^2-^; detection limit 5 µg/L) and Fe (II; detection limit 0.01 mg/L) were determined in the field with a portable Hach spectrophotometer (model DR2800) and Hach reagents (Hach Company, Loveland, CO, USA) following established protocols (Colman et al., 2016). Sulfide and Fe (II) concentrations were measured on water samples collected by a high-density polyethylene sampling staff (turbid water samples were filtered through pre-sterilized 0.22 µm Sterivex filters (EMD Millipore, Billerica, MA, USA)), while all other parameters were measured *in situ* near the site of sediment collection. Sediments were collected into sterile conical centrifuge tubes using the sampling staff, placed on dry ice in the field, then transferred to storage at –80°C °until further processing.

**Figure 2.**
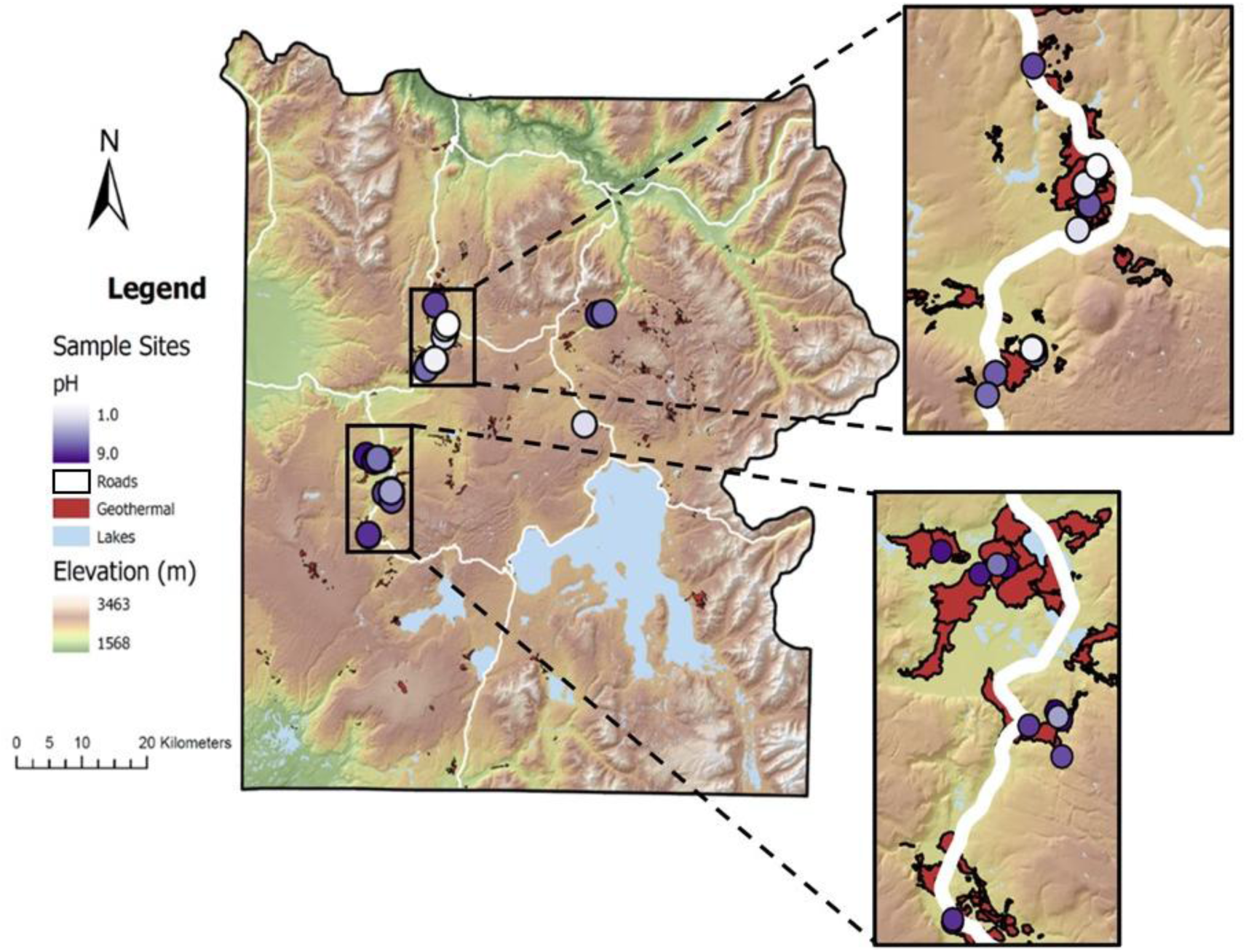
Map of sampling locations colored by pH in Yellowstone National Park, USA 2018 to 2022. Major roads are shown in white and geothermal areas are shaded in red.

### Lipid Extraction and Preparation

Sediment samples were freeze-dried and stored at room temperature before extraction. Total lipid extracts (TLEs) were obtained by modified Bligh-Dyer extraction (Bligh and Dyer, 1959; Weber et al., 2017). Sediments (∼1 g) were sonicated in 2:1:0.8 methanol:dichloromethane:trichloroacetic acid (MeOH:DCM:TCA) buffer (TCA, 50 g/L, pH = 2.1), centrifuged, and phase-extracted in four rounds. Organic phases underwent a final water rinse before concentration under N_2_ gas and reconstitution in 2:1 DCM:methanol. Where necessary, multiple samples were merged for larger lipid yields (Table S1). TLEs were treated with activated (1 M HCl, rinsed with H_2_O, MeOH, DCM, and hexane) copper at room temperature overnight to remove S^0^ (Boyd et al., 2011). TLEs were separated into core lipid (CL) and intact polar lipid (IPL) fractions by solvent elution through SiO_2_ columns as described in Pitcher et al. (2009) and stored in DCM at –20°C. The polar headgroups of IPL fractions were cleaved (5% HCl in MeOH, 3 hr). IPL-derived lipids were brought to a pH of 4 to 5 using 1 M KOH in MeOH, followed by four rounds of liquid-liquid extraction. CL– and IPL-derived lipid fractions were reconstituted in a mixture of 99:1 hexane:isopropyl alcohol (IPA) with addition of 100 ng of C_46_-GTGT internal standard (Huguet et al., 2006). All samples were filtered through 0.45 µm PTFE filters (4 mm diameter) before analysis.

### GDGT Identification and Relative Quantification

All iGDGT fractions were analyzed by ultra-high performance liquid chromatography – atmospheric pressure chemical ionization – mass spectrometry (UHPLC-ACPI-MS) using an Agilent 1290 Infinity series UHPLC system coupled to an Agilent 6460 triple-quadrupole mass spectrometer (QQQ MS), consistent with methods used in earlier work (Zhou et al., 2020; Blewett et al., 2020). GDGTs were separated by injecting 10 µL of sample onto two coupled ACQUITY UPLC BEH Amide Columns (1.7 µm, 2.1 x 150 mm) held at 50°C (Becker et al., 2013) with a constant solvent flow rate (0.5 mL/min) and solvent mixtures A (pure hexane), B (90:10 hexane:IPA), and D (30:70 MeOH:IPA). The program started at 98:2 A:B with a linear gradient to 3% B by 4 minutes, 10% B by 10 minutes, 20% B by 20 minutes, 50% B by 35 minutes, and 100% B by 40 minutes followed by a one-minute hold. At 41.01 minutes, the program was set to 70:30 B:D, ramping to 100:0 B:D by 46 minutes and 98:2 A:B by 47 minutes, holding until the total run time of 65 minutes. The QQQ-MS was operated in single ion monitoring (SIM) mode with a dwell time of 128 ms and fragmentor voltage of 75 V. GDGT relative abundances were determined by manual integration of ion chromatograms with mass to charge ratios (*m*/*z*) starting at 1302.3 for iGDGT-0 and decreasing by increments of 2 down to 1286.3 *m*/*z* for iGDGT-8.

### Data Reduction and Statistical Analyses

Two RI values were calculated: with and without crenarchaeol (Eq. 1 and 2). In the numerator, the relative abundance of each lipid is multiplied by the number of cyclopentyl rings in the lipid structure (Taylor et al., 2013; Zhang et al., 2016). For this reason, crenarchaeol is incorporated as a four-ringed member in Equation 1 because the fifth ring is the enigmatic cyclohexyl ring whose contribution to membrane dynamics is so far unconstrained (Zhang et al., 2016). See the Supplement for a discussion of the merits and drawbacks of including crenarchaeol as a four-ringed contributor to RI; we conclude that it is appropriate to exclude crenarchaeol from RI calculations if the intended purpose is to investigate membrane permeability.

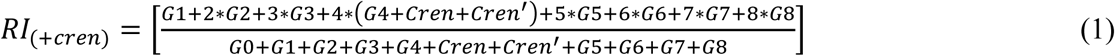

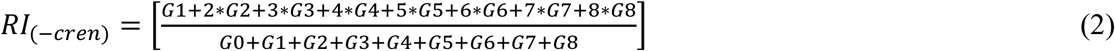

Given that neutral pH varies with temperature, we calculated the temperature-dependent neutral pH, i.e., the neutrality line, from 0 to 100°C (Eq. 3; Justnes et al., 2020).

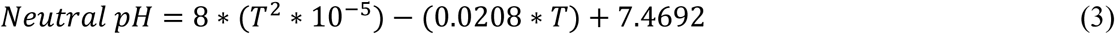

Statistical analyses were performed using R Statistical Software (v4.4.0; R Core Team, 2024). Scripts for statistical runs and plots are available at https://zenodo.org/doi/10.5281/zenodo.12603497. Quadratic and interaction terms were tested in modelling, but these were not substantially more powerful than reported models. All input variables of regression models are normalized using the base *scale*() function of R to prevent skew due to parameter magnitude.

## 3.0 Results

### 3.1 Lipid characteristics among YNP hydrothermal springs

Geochemical parameters for the YNP samples collected in this study (N = 41; Supplement Table S1) include temperature (28 to 93°C), pH (1.1 to 9.0), dissolved oxygen (DO; 1.6 x 10^-7^ to 6.2 x 10^-5^ M), specific conductivity (SPC; 50 to 5380 µS/cm), oxidation reduction potential (ORP; –330 to 330 mV), dissolved Fe (II) (B.D. to 3.5 x 10^-5^ M) and dissolved sulfide (S^2-^; B.D. to 2.5 x 10^-4^ M) concentrations. None of these parameters were normally distributed, so Spearman’s rho correlations were used to evaluate their relationships with lipid characteristics. The relationships between these environmental variables, crenarchaeol relative abundance, and Ring Index (see Table S1 for individual site values) were calculated using simple (SLR) and multiple linear regressions (MLR) of all possible geochemical parameter combinations for both CL and IPL fractions. Model results reported below are for CL fractions and statistical results for both fractions are compared in the Supplemental Information (Tables S2, S3, S5, S6), as CL lipids are often more abundant and represent a more integrative record of iGDGT production than IPL lipids. The RI results reported here include crenarchaeol as a four-ringed member, as seen in previous studies (Zhang et al., 2016).

The non-linear Spearman’s rho tests demonstrated that pH and Fe (II) concentration were the only variables that were significantly associated with CL and IPL crenarchaeol relative abundance for the 41 new YNP samples (Table S2). However, significant relationships between Fe (II) concentration and crenarchaeol relative abundance are likely due to the strong correlation of Fe (II) with pH (Figure S6; Amenabar and Boyd, 2018), with the latter being the likely true driver of crenarchaeol relative abundance. None of the individual environmental variables exhibited significant (*p* < 0.05) simple linear regressions with crenarchaeol relative abundance, but several combinations of variables exhibit significant multiple linear regression relationships with crenarchaeol (Table S3). Of all possible variable combinations (127 models), the most important environmental parameters for predicting crenarchaeol relative abundance were pH, SPC, and ORP. All five multiple linear regression models with the highest adjusted R^2^ values include these three variables, which alone explain 19% of the variance (adj-R^2^ = 0.19) (Table S4). The model that incorporates sulfide as a fourth variable in addition to pH, SPC, and ORP is the best overall predictor (adj-R^2^ = 0.21). In contrast, including DO or temperature as the fourth variable decreases the model explanatory power (adj-R^2^ = 0.17).

Spearman’s rho tests of RI values showed that pH, ORP, and Fe (II) concentration were significantly associated with both the CL and IPL fractions from the YNP thermal spring samples (Table S5). Only the CL fraction had a non-linear association with temperature. Simple linear regression results matched those of the Spearman’s rho correlation, with pH, ORP, and Fe (II) concentration correlating with both CL and IPL fractions, while temperature only correlated with the CL fraction (Table S6). Sulfide has a significant linear relationship with IPL-RI, but there is no non-linear association (Spearman’s) between these two variables. Of all 127 possible multiple linear regression models of RI, the ten most significant (*p* < 0.05) models include pH and DO, while Fe (II) is present in eight, temperature in six, sulfide in five, SPC in three, and ORP in two (Table S7). The best explanatory models incorporate pH, temperature, DO, and Fe (II) (adj-R^2^ = 0.80), or just the three variables pH, DO, and Fe (II) (adj-R^2^ = 0.79). The addition of a fourth variable other than temperature slightly decreases model predictive power, but all ten most significant models exhibit an adjusted R^2^ range of 0.79 to 0.80. In high temperature hot springs that exclude photosynthesis, the availability of DO is controlled, at first order, by temperature-dependent O_2_ solubility (Shock et al., 2010), indicating that temperature may be the real driver of crenarchaeol relative abundance in cases above in which DO is implicated as an important variable.

### 3.2 Comparison to previously published studies

We compiled published thermal spring lipid samples (N = 299 including 41 from this study) that report pH, temperature, and iGDGT abundances, specifically including crenarchaeol abundance (Pearson et al., 2004; Zhang et al., 2006; Schouten et al., 2007; Pearson et al., 2008; Pitcher et al., 2009; Zhao et al., 2011; Burgess et al., 2012; He et al., 2012; Li et al., 2013; Boyd et al., 2013; Wu et al., 2013; Paraiso et al., 2013; Jia et al., 2014; Xie et al., 2015). We include relevant culture data in Figure 3 for context but exclude this data from our statistical analyses to focus on environmental samples. For previous studies that split iGDGTs into CL and IPL fractions, the CL crenarchaeol relative abundances were used in this compilation since CL-iGDGTs are generally dominant and represent a more integrative record of iGDGT production than IPL lipids. The compiled dataset affords greater environmental context than the dataset from YNP samples alone; however, it reduces the available environmental variables to pH and temperature, which we focus on in our interpretation.

**Figure 3.**
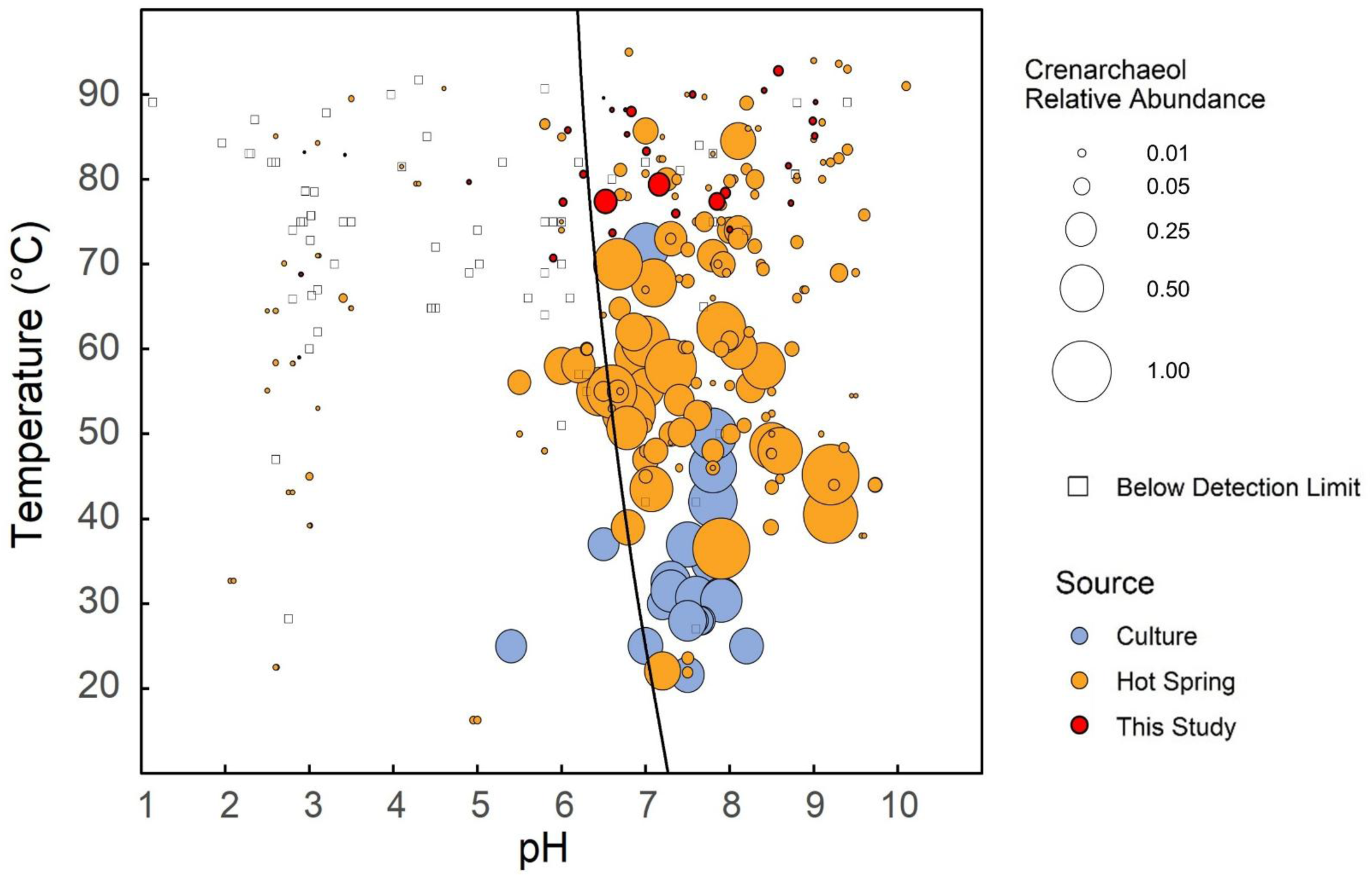
Relative abundances of crenarchaeol for all compiled thermal spring samples (N = 299) and culture samples (N = 33). Samples with no detectable crenarchaeol are represented by squares and those with crenarchaeol present are circles filled by the source of the data. Area of circles correlates with crenarchaeol relative abundance. The black curve is the temperature-dependent neutral pH line (Equation 3).

Across all sites, crenarchaeol relative abundances were highest to the alkaline side (≥ 7.0 pH) of the temperature-dependent neutral pH line (Equation 3), while acidic (< 7.0 pH) springs frequently have non-detectable levels of crenarchaeol (Figure 3). The 109 acidic samples have an average fractional crenarchaeol abundance of 0.02 (2% of all iGDGTs or CL iGDGTs), while that of the 190 alkaline samples is 0.10, with many examples > 0.50. Crenarchaeol was undetectable in 35/109 (32.1%) of acidic samples vs. 12/190 (6.3%) of alkaline samples. Crenarchaeol relative abundance also shows a weak relationship to temperature. Of 201 samples from environments ≥ 60°C, 79.6% have detectable crenarchaeol. Ninety-eight samples are from < 60°C, of which 93.9% contain detectable crenarchaeol. Additionally, at ≥ 60°C, the average crenarchaeol fractional abundance is 0.04 (0.05 without zero values) and < 60°C, the average is 0.14 (0.15 without zeroes).

### 3.3 Statistical analyses

Simple and multiple linear regression models were conducted to evaluate the influence of pH, temperature, and the combination of both variables for the full dataset (Table 1). MLR models incorporating interactions between temperature and pH and quadratic variables were run, but did not noticeably improve R^2^ values (0.13 with interactions and 0.15 with quadratic variables for the crenarchaeol models), so linear models are shown for simplicity. Non-linear Spearman’s rho correlations indicated that crenarchaeol is strongly associated with both pH and temperature (*p* = 1 x 10^-13^ and 5 x 10^-9^, respectively), while Ring Index (Equation 1) is associated with pH (*p* = 7 x 10^-9^) but not with temperature (*p* = 0.8). While all R^2^ values show relatively low explanatory power, pH dominates predictions of RI, while both pH and temperature are significant individual predictors of crenarchaeol relative abundance.

**Table 1.** Outputs of simple and multiple linear regression models for pH and temperature relationships with crenarchaeol relative abundance and Ring Index of all compiled data (N = 299). Adjusted R^2^ values are reported, and significant p-values are indicated by bold italics. Note that negative R^2^ values indicate a fit worse than a horizontal line.

|  | Parameter(s) | Crenarchaeol |  | RI |  |
| --- | --- | --- | --- | --- | --- |
| | | p-value | adjusted $R^2$ | p-value | adjusted $R^2$ |
| Simple Linear Regression | pH | **<0.001** | 0.033 | **<0.0001** | 0.15 |
| | Temp ($^{\circ}\text{C}$) | **<0.0001** | 0.073 | 0.62 | -0.0025 |
| Multiple Linear Regression | pH + Temp ($^{\circ}\text{C}$) | **<0.0001** | 0.11 | **<0.0001** | 0.15 |

The two variables are visualized in Figure 4. Relative crenarchaeol abundance is lower in many high-temperature samples and lower or absent in most of the lowest-pH samples (Figures 4A, 4B). RI has no apparent patterns associated with temperature, but high RI values are often identified in low-pH samples (Figures 4C, 4D). While the linear regression models are statistically significant, all these correlations are modest (Table 1), and some of the highest RI values occur at high pH values (e.g., Figure 4C).

**Figure 4.**
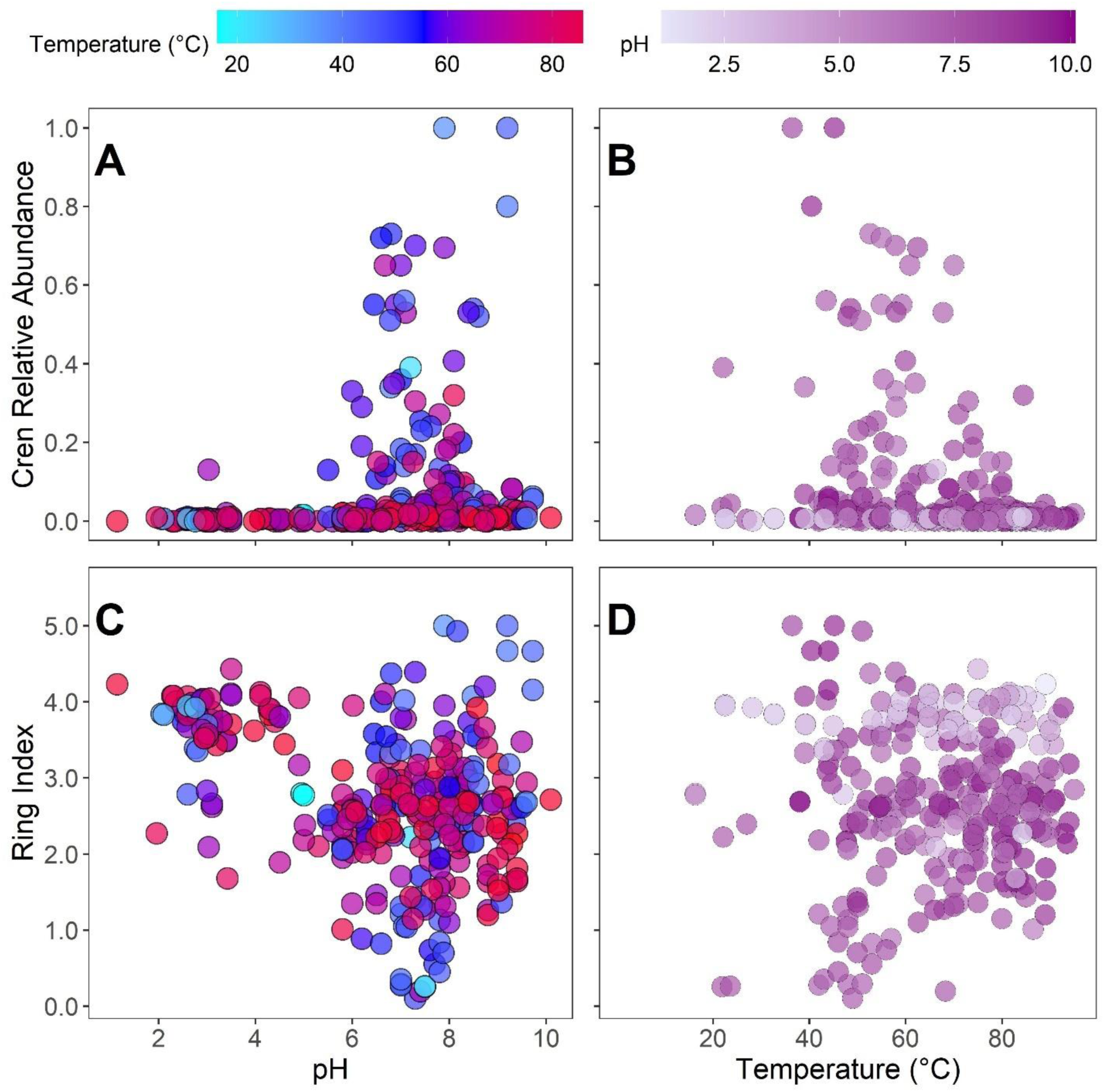
Crenarchaeol relative abundance versus (A) pH and (B) temperature for the 299 compiled samples, with fill by the other variable. Ring Index versus (C) pH and (D) temperature for the 299 compiled samples, with fill by the other variable.

## 4. Discussion

### 4.1 Environmental Predictors of Ring Index

Studies of marine systems generally consider temperature to be the primary driver for iGDGT cyclization (Schouten et al., 2002; Kim et al., 2008; Kim et al., 2010; Tierney, 2012), which may result from the relative geochemical uniformity of modern open-ocean environments. In marine waters, deviations from global average RI or TEX_86_ vs. temperature correlations are associated with pH, DO concentration, ammonia oxidation rate, growth phase, and salinity (Qin et al., 2015; Elling et al., 2015; Hurley et al., 2016; Hurley et al., 2018), demonstrating that variables other than temperature may be of greater importance in other settings (Boyd et al., 2011; Elling et al., 2015; Cobban et al., 2020). It is likely that control of iGDGT cyclization in thermal springs is different from the marine system, because thermal springs generally span larger gradients of environmental variables, including but not limited to pH. Previous studies examining iGDGT cyclization in thermal springs have yielded conflicting conclusions. Temperature, pH, salinity, redox potential, bicarbonate concentration, growth rate, and DO all have been reported to correlate with iGDGT cyclization (Pearson et al., 2004; Boyd et al., 2013; Elling et al., 2015; Qin et al., 2015; Feyhl-Buska et al., 2016; Evans et al., 2018; Cobban et al., 2020; Zhou et al., 2020). While the degree of importance of each of these variables on iGDGT cyclization has not been definitively established, some studies conclude that pH has a greater influence than temperature. For example, thermal springs spanning pH 5.5 to 7.2 in New Zealand showed increased RI at higher temperatures, but more acidic springs (pH 2.1 to 5.5) did not demonstrate this temperature effect (Kaur et al., 2015). These results indicate that acidity is more important than temperature for affecting archaeal membrane composition, potentially in direct response to pH-induced stress. The discrepancies among studies may be due to sampling relatively narrow ranges of geochemical and geophysical parameters, or it may result from the competing effects of several environmental factors (e.g., temperature and pH; Pearson et al., 2008; Wu et al., 2013; Boyd et al., 2013; Xie et al., 2015). We synthesize prior works and our new observations from YNP to consider the environmental stressors of pH and temperature together, rather than in isolation.

Here, our analysis of compiled global thermal springs (N = 299) demonstrates that pH significantly influences RI, while temperature does not. Linear regression models for the 41 YNP samples indicate that pH alone predicts a majority of the RI distribution in thermal springs (CL, 64%; IPL, 55%), consistent with previous thermal springs studies that concluded pH is the primary predictor of both RI (Pearson et al., 2008; Boyd et al., 2013) and the distribution of genes related to cyclopentyl ring production (Blum et al., 2022). ORP emerges as an important predictor of RI in linear regressions of the YNP dataset (CL = 31%; IPL = 31%), consistent with earlier findings that redox conditions are important for archaeal lipid stress responses (Cobban et al., 2020). Spearman’s rho correlations and simple linear regression models highlight the influence of ORP on membrane lipid cyclization. However, ORP has a direct relationship with pH (James and Lytle, 2004), which likely contributes to its association and simple regression significance with RI. Similarly, while DO may indicate redox conditions, its temperature-dependent solubility confounds interpretation of an independent redox state signal. Following ORP (or DO), Fe (II) emerges as a potential control on archaeal membrane GDGT composition (predictive power: CL, 22%; IPL, 22%), but the covariation of Fe (II) with decreasing pH makes it difficult to determine whether Fe (II) exerts any independent effect (Figure S6). Sulfide concentration exhibits minimal predictive power, with influence only on IPL-RI in a linear regression (11%), but not on CL-RI, nor any significant Spearman’s rho association. As sulfide is a soluble chemical species influenced by pH and redox state, any significant association with iGDGT cyclization may be a result of covariation with other variables. Temperature is associated with CL-RI but not IPL-RI, which may be explained by differing CL and IPL crenarchaeol relative abundances in one or two samples that drive the CL correlation over the significance threshold (see Supplement for detailed discussion). It is also possible that temperature influences RI over long time periods (CL), while other variables have greater control over ring abundances over shorter time frames (IPL). In other words, IPL composition is more responsive to recent geochemical fluctuations that may not have been captured in this latitudinal study design.

### 4.2 Predictors of Crenarchaeol Relative Abundance

Crenarchaeol relative abundance, like RI, was associated with multiple environmental variables. While pH, temperature, and redox explain the majority of the variance of RI (MLR: R^2^ = 0.80), temperature and pH are both only weakly associated with crenarchaeol relative abundance (SLR: Temp R^2^ = 0.07, pH R^2^ = 0.03) and combining them adds minimal explanatory power (MLR: R^2^ = 0.11; See Table 1). The variables examined in this study do not sufficiently explain cyclohexyl ring distributions, and simple linear regressions cannot determine the rank order of influence of pH and temperature on crenarchaeol distributions. The relative importance of the two variables also is inconclusive for the non-linear Spearman’s rho tests. The clear tradeoff between statistical power (N = 299 and N = 41) and geochemical detail (two variables versus seven) between the compiled and the Yellowstone-only datasets demonstrates the need for consistent variable collection and robust analyses to evaluate a ranked list of variables important for crenarchaeol production.

While temperature is associated with crenarchaeol relative abundance in the global thermal spring dataset, it is not associated with crenarchaeol relative abundance in the more geochemically detailed YNP dataset. The latter observation may be attributed to a lower representation of samples below 60°C for the newly collected YNP dataset (N = 2) compared to the global dataset (N = 96). This result suggests that temperature is associated with crenarchaeol relative abundance, but that its relationship is weaker than that of pH. Variables present exclusively in the Yellowstone dataset (e.g., ORP and DO) covary with pH and temperature, and may represent covariation signals. An early examination of crenarchaeol from thermal springs found that bicarbonate concentration (which is partially controlled by the influence of pH on DIC speciation) and not temperature correlated with crenarchaeol distribution (Pearson et al., 2004). While that study measured pH, samples were limited to six regional springs that spanned pH values of 6.4 to 9.2, a range that would not show the pH signal observed in our current study. A similarly confirmatory study examined 27 YNP hot spring samples and demonstrated that crenarchaeol relative abundance correlates with hot spring chemistry, not temperature (Boyd et al., 2013). CL crenarchaeol and crenarchaeol isomer relative abundances correlated with pH and NO_2_^-^ concentration, while CL crenarchaeol relative abundance demonstrated an inverse correlation with NH_4_^+^ concentration, consistent with AOA production of crenarchaeol in these springs (Boyd et al., 2013). In IPL fractions, crenarchaeol relative abundance correlated with Cl^-^ concentration (indicative of water source) while crenarchaeol isomer relative abundance correlated with Fe (II) concentration, which is controlled by pH (Boyd et al., 2013). These results further indicate that hydrothermal water chemistry influences crenarchaeol relative abundance in archaeal lipidomes.

The strong association between pH and crenarchaeol relative abundance is perhaps unsurprising. Without protective adaptations to external pH, a cell’s pH homeostasis can be disrupted, thus slowing enzyme activity, destabilizing other proteins and nucleic acids, and potentially resulting in cell death (Slonczewski et al., 2009). As most literature on iGDGT distributions is focused on marine paleoclimate applications, the influence of pH on crenarchaeol abundance has remained uncharacterized due to the narrow range of pH observed in the marine realm. The average pH of Earths’ surface oceans is 8.07 ± 0.02 between 60° North and 60° South (Jiang et al., 2019), but *Nitrososphaeria* also thrive in less alkaline marine thermoclines (pH 7.5 to 7.2 from 200 to 600 m depth; Clayton and Byrne 1993; Palmer et al., 1998; Church et al., 2010). The wide pH range of terrestrial thermal springs (1.14 to 10.10) covered in this compilation shows that crenarchaeol is present over a broad range of pH values (1.96 to 10.1) but is most abundant (> 30% of core/total iGDGTs) from pH 6.0 to 9.2. While a pH range from 6.0 to 9.2 is relatively narrow for thermal springs, it is much wider than that of the modern surface ocean (8.0 to 8.25; Jiang et al., 2019).

Differences in parameters other than pH that contribute to multiple linear regression models of crenarchaeol and RI may indicate that nuanced or independent processes influence production of cyclohexyl rings in iGDGTs or may reflect co-variation of soluble chemical parameters with pH and temperature. In addition to pH, DO and Fe (II) are statistically important predictors of RI, while those of crenarchaeol relative abundance are SPC, ORP, and temperature. While these secondary variables could cause membrane stress that would alter lipid production (Cobban et al., 2020; Zhou et al., 2020), it may be more likely that these weak correlations of soluble geochemical parameters with lipid compositions are due to co-variation with parameters such as pH and temperature. While cyclohexyl rings could have a specific relationship to cellular redox conditions, this is not supported by culture experiments that have altered redox state and observed no significant change in crenarchaeol production (Qin et al., 2015).

### 4.3 Temperature and pH Optima for Crenarchaeol Production

To test the hypothesis that crenarchaeol may optimize thermophilic membranes for more mesophilic settings, the optimal temperature and pH for crenarchaeol relative abundance were calculated (total data, N = 299; Figure 5). The pH range (discussed hereafter as bins shown in Figure 5) with the highest mean crenarchaeol relative abundance is 7.0 to 7.5 while bins from pH values of 6.5 to 8.5 also have high sample counts with elevated means and upper quartiles relative to those outside of this pH range (Figure 5A). The bin from pH 9.5 to 10 also has high crenarchaeol relative abundance (0.01 to 0.06), but this may be an artifact of the small site number (N = 5), demonstrating the need for further quantification of archaeal lipids in alkaline springs. The temperature bin with the highest mean crenarchaeol relative abundance spans 50 to 55°C, while bins from 40 to 60°C have elevated mean and upper quartile values relative to those outside this range (Figure 5B). In general, bins from 40 to 90°C have high sample counts while bins from 15 to 40°C and above 90°C have low sample counts, explaining the high upper quartile value of the bin from 35 to 40°C (N = 7).

**Figure 5:**
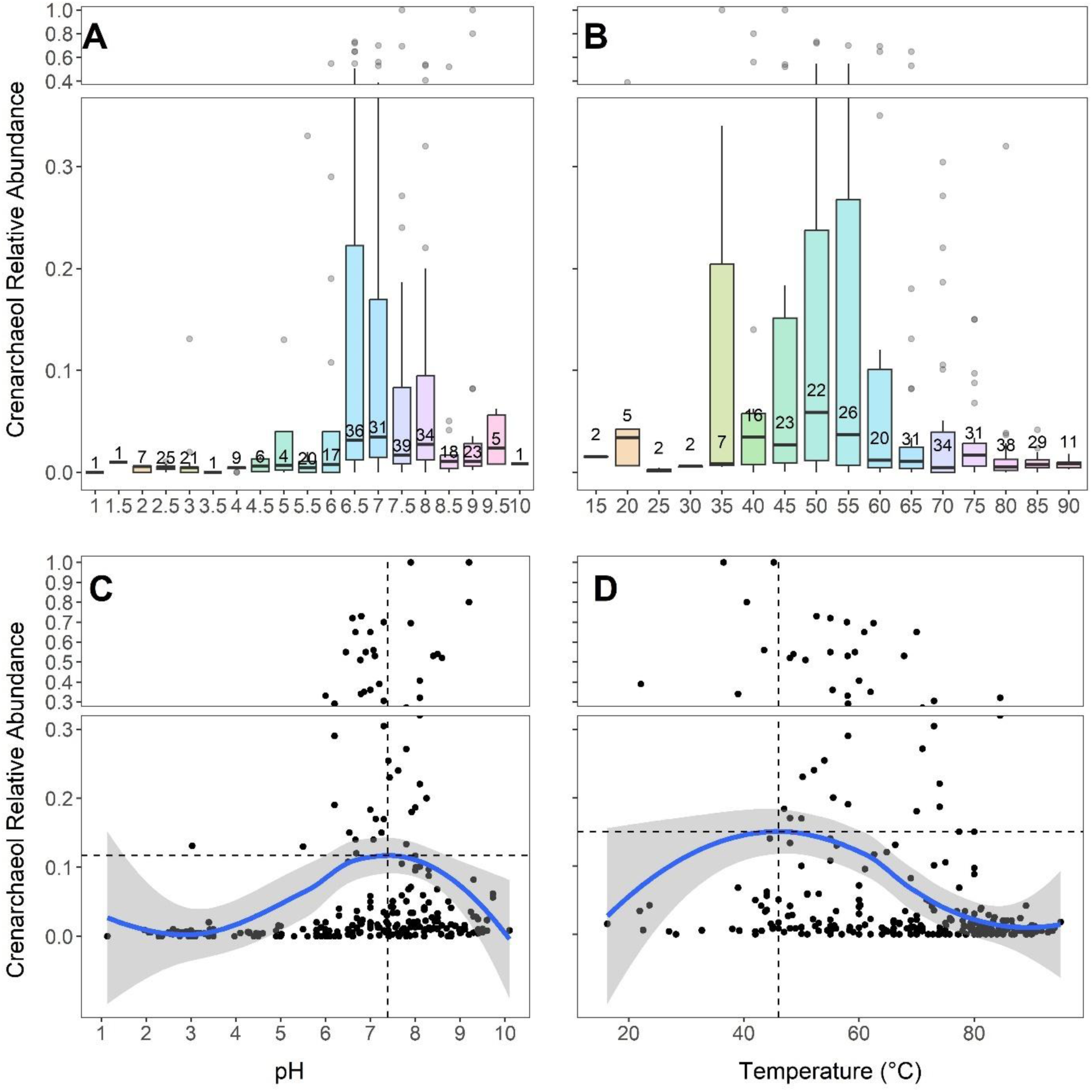
Boxplots (A &B) and LOESS smoothing curves (C&D) of crenarchaeol relative abundances of compiled data (N = 299). Data ranges are binned by 0.5 pH units (A) and 5°C temperature units (B) with bins labelled by their minimum bound. Median and quartile abundances are visualized in boxes and maximum and minimum values are indicated by whiskers. Sample counts for each bin are included above mean value indicators in A and B. The functions plotted in C and D have 95% confidence intervals shaded in gray. Dashed lines in C and D represent x and y values where crenarchaeol reaches its maximum. Y-axis breaks and scale changes are utilized to allow outliers to be visualized.

The LOESS (locally weighted smoothing; Figures 5C and 5D) functions show the optimal environmental pH for maximizing the relative abundance of crenarchaeol is 7.4 (corresponding to 0.12 LOESS-predicted relative abundance), consistent with the boxplot having the highest mean (pH 7 to 7.5) in Panel 5A. The optimal temperature for maximizing the relative abundance of crenarchaeol is 46°C, (0.15 LOESS-predicted relative abundance). While this value is not within the highest-mean bin of Panel 5B (50 to 55°C), it does fall within the broad grouping of temperatures showing elevated mean crenarchaeol relative abundances (40 to 60°C); this is consistent with temperature having a weaker overall control on iGDGT, and specifically crenarchaeol, patterns compared to pH. Scatter in Panel 5D shows that crenarchaeol can also be found in high relative abundances above 80°C. Note that thermal spring data are expected to have some degree of noise due to environmental variability over several timescales including changes in volcanic/hydrothermal activity, seasonality, and diurnal cycles (Payne et al., 2019; Colman et al., 2021; see Supplement for further discussion).

The analysis shown in Figure 5 corroborates the observed circumneutral pH threshold identified in Figure 3, in which instances of abundant crenarchaeol track the temperature-dependent neutral pH line. The directionality of the relationship of crenarchaeol with environmental parameters is consistent with previous hypotheses that crenarchaeol acts to expand archaeal membranes (in opposition to cyclopentyl rings; Sinninghe Damsté et al., 2002). Acidic environments have an excess of protons, promoting the need for cell membrane compaction through cyclopentyl ring production to prevent energetic imbalances (Boyd et al., 2011). If crenarchaeol expands membrane fluidity, crenarchaeol concentrations should be expected to be lower in acidic environments. While there exists an abrupt acidity cutoff for high relative crenarchaeol production, there is no apparent upper pH limit (Figure 3). This may indicate that high pH is not as stressful as acidic pH for crenarchaeol-producing archaeal cells. Nevertheless, further study of alkaline environments from pH 9 to 12 could illuminate a decrease in crenarchaeol production if an excess of hydroxyl ions poses significant stress to archaeal cells. Another consideration when interpreting the pH optimum of crenarchaeol is the pH-dependent speciation of soluble metabolic substrates, such as NH_3_ utilized during nitrification by AOA. While the pKa of the NH_4_^+^-NH_3_ system is strongly temperature dependent (Amend and Shock, 2001), NH_3_ is scarce in acidic settings, potentially decreasing crenarchaeol production by AOA that lack metabolic substrates in acidic systems. However, the production of crenarchaeol in environments in which AOA are rare (>75°C, acidic pH) warrants further explanation, as discussed below in section 4.5.

### 4.4 Recommendations for Future Studies

Given the association of crenarchaeol relative abundance with multiple environmental parameters identified in this study, comprehensive analyses that quantify the relative influence of environmental variables on crenarchaeol relative abundance will help guide interpretations of this biomarker in the sediment record. Future work manipulating individual variables, particularly pH, in culture would help strengthen, qualify, or contradict the conclusions from our environmental dataset. Additionally, our compiled global literature datasets all have associated thermal temperature and pH measurements, but other parameters, such as ORP, SPC, DO, and various dissolved ions, are inconsistently collected across studies, thus decreasing statistical power. Future studies investigating iGDGT cyclization should collect a standardized suite of environmental parameters to tease apart the relative influence of multiple factors. Repeated time-series sampling within geochemically dynamic springs may also provide valuable insight into the response of archaeal lipid membrane compositions to short-term geochemical variability.

An important knowledge gap in the crenarchaeol literature is the identity of the gene(s) encoding cyclohexyl ring formation (Zeng et al., 2022). Laboratory experiments aiming to produce abundant crenarchaeol from enrichment cultures will benefit from targeting temperatures from 40 to 55°C and pH values from 6.5 to 8.0 (Figures 5A, 5B), while pairing comparative transcriptomics or proteomics data to identify putative cyclohexyl ring biosynthesis enzyme(s). To test the long-standing belief that crenarchaeol is specific to *Nitrososphaeria* (Pearson and Ingalls, 2013; Schouten et al., 2013), correlating the relative abundance of crenarchaeol with different taxonomic groups could either support or contradict this assertion (Sinninghe-Damsté et al., 2018).

### 4.5 Explanations for Crenarchaeol Optima in Natural Hydrothermal Springs

#### Environmental Distributions of Crenarchaeol Producers

A straightforward interpretation of the environmental crenarchaeol optima may be that the *Nitrososphaeria* that produce crenarchaeol grow optimally in circumneutral and mesophilic settings, as several cultivation studies show that AOA are most abundant below ∼75°C and at or above neutral pH (Könneke et al., 2005; de la Torre et al., 2008; Hatzenpichler et al., 2008; Elling et al., 2015). This interpretation is supported by the correlation of *amoA* gene abundance with the distribution of crenarchaeol in 27 thermal spring samples (Boyd et al., 2013). However, AOA are absent or scarce in hyperthermal (>75°C) springs (Hou et al., 2013; Podar et al., 2020; Colman et al., 2024), indicating that other groups may be producing crenarchaeol at high temperatures, and potentially more moderate spring conditions. Not all *Nitrososphaeria* are AOA, and novel groups have been shown to be thermoacidophilic (Beam et al., 2014; Kato et al., 2019). The non-AOA group *Conexivisphaerales* is abundant in YNP hydrothermal springs below pH 5 (Supplementary Figure S2; analysis of the *Nitrososphaeria* metagenome-assembled genome (MAG) data from Colman et al., 2024). The AOA groups demonstrate the expected preference for circumneutral to alkaline settings, while the non-AOA groups inhabit either acidic or circumneutral pH ranges (Colman et al., 2024). So far, one thermoacidophilic isolate exists from the non-AOA *Nitrososphaeria* (*Conexivisphaera calidus*; Kato et al., 2019), but its lipidome remains uncharacterized. Therefore, it remains unknown whether the acidophilic *Nitrososphaeria* produce crenarchaeol, but the presence of only trace amounts of crenarchaeol below pH 5 indicate that crenarchaeol does not constitute a large portion of the membrane in *Conexivisphaerales* or other *Nitrososphaeria* from lower pH systems. In contrast, a non-AOA *Nitrososphaeria* group, the *Caldarchaeales,* preferentially inhabit circumneutral settings, and are present in hyperthermal springs (Colman et al., 2024). While a tungsten-dependent representative of the *Caldarchaeales* (*Wolframiiraptor gerlachensis*) has been cultivated in a stable enrichment culture (Buessecker et al., 2022), the lipidomes and potential for crenarchaeol production of *Caldarchaeales* members remain uncharacterized. Future work to characterize the membrane composition of non-AOA *Nitrososphaeria* isolates and to identify the gene encoding the enzyme for cyclohexyl ring formation will be critical to constrain the phylogenetic distribution of crenarchaeol production and will allow us to better differentiate phylogenetic versus geochemical controls on crenarchaeol distributions in nature and throughout the geologic record.

#### A Membrane-Expanding Function

The circumneutral and mesophilic crenarchaeol optima estimated in this study may also be interpreted to be consistent with the hypothesized function of crenarchaeol to decrease membrane lipid packing. AOA are thought to have radiated from a thermophilic ancestor (Abby et al., 2020; Luo et al., 2024), which would have produced lipid membranes with high cyclopentyl ring abundances to protect against extremes in temperature. Upon expansion into cooler marine waters, these organisms would have required a mechanism to increase membrane fluidity to adapt to less “stressful” environments. While our data cannot determine whether crenarchaeol facilitated this evolutionary transition, our environmental optima and the thermophilic nature of AOA precursors agree with the hypothesized membrane-expanding function of crenarchaeol. Our analysis spans a larger range of environmental variables and has a higher sampling density than previous studies, indicating that the temperature optimum of crenarchaeol from this study (46°C) is likely representative, while the prior estimated optimum of 40°C (Zhang et al., 2006) is within the elevated range of the 95% confidence interval of the model (Figure 5). Our temperature optimum agrees with the respective 30-65°C (Robert and Chaussidon, 2006) and 47°C (Grossman and Joachimski, 2022) sea-surface temperature reconstructions during the predicted dates of the transition of AOA into the ocean (1017 Ma, Ren et al., 2019; 509 Ma, Yang et al., 2021) and the 33 to 42°C range (Bice et al., 2006) during the proposed expansion of non-thermophilic marine *Nitrososphaeria* during the mid-Cretaceous anoxic event (ca. 112 Ma). Future studies examining the distribution of crenarchaeol in the geologic record and identifying the gene encoding the enzymes for cyclohexyl ring formation will allow determination of the potential function of crenarchaeol in the evolutionary history of archaea. The current study provides the environmental context for mechanistic investigations into the distribution of crenarchaeol in geochemically diverse settings.

## Acknowledgements

This work was supported by a Dartmouth Stamps Scholarship (ANC), the Walter and Constance Burke Award of Dartmouth College (WDL), an NSF-EAR grant (WDL), NASA grant 80NSSC19M0150 (DRC and ESB), and an ACS-PRF award (AP). We thank C. Hendrix, S. Gunther, and A. Carlson at Yellowstone National Park for permitting research access under YNP permit #YELL-05544. We also wish to thank M. Palucis (Dartmouth) for help with the design of the Yellowstone map. We thank D. Payne (MSU), L. Denoncourt (MSU), and John Spear (Colorado School of Mines) for field assistance.

## Supplemental Information

### Figures

**Figure S1.**
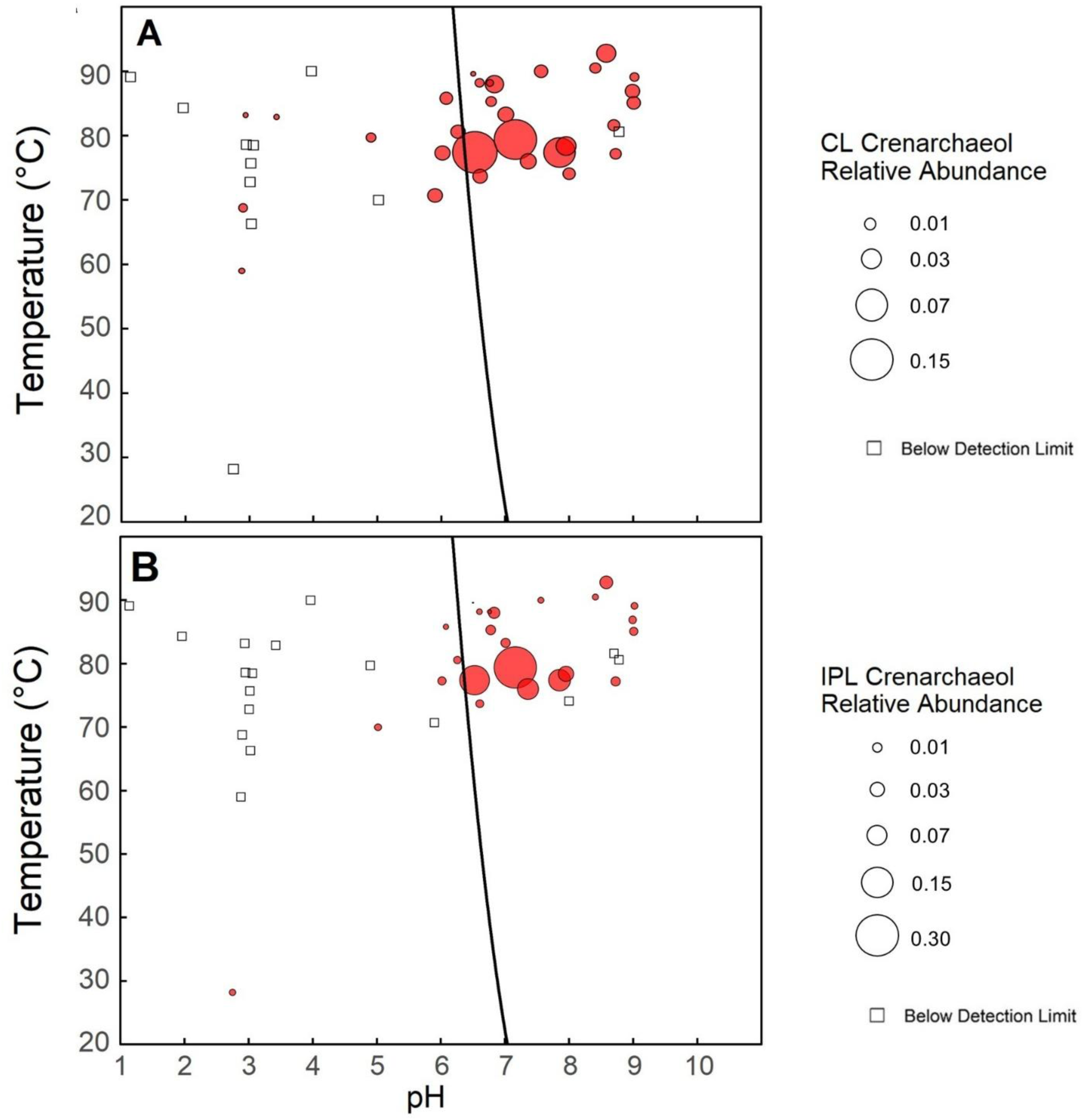
Relative abundances of (A) CL and (B) IPL crenarchaeol in temperature-pH space for 41 Yellowstone field samples collected from 2018-2022. Bubble area is proportional to relative abundance of crenarchaeol while squares represent non-detectable abundances. The black curve is the temperature-dependent neutral pH line (Equation 3).

**Figure S2.**
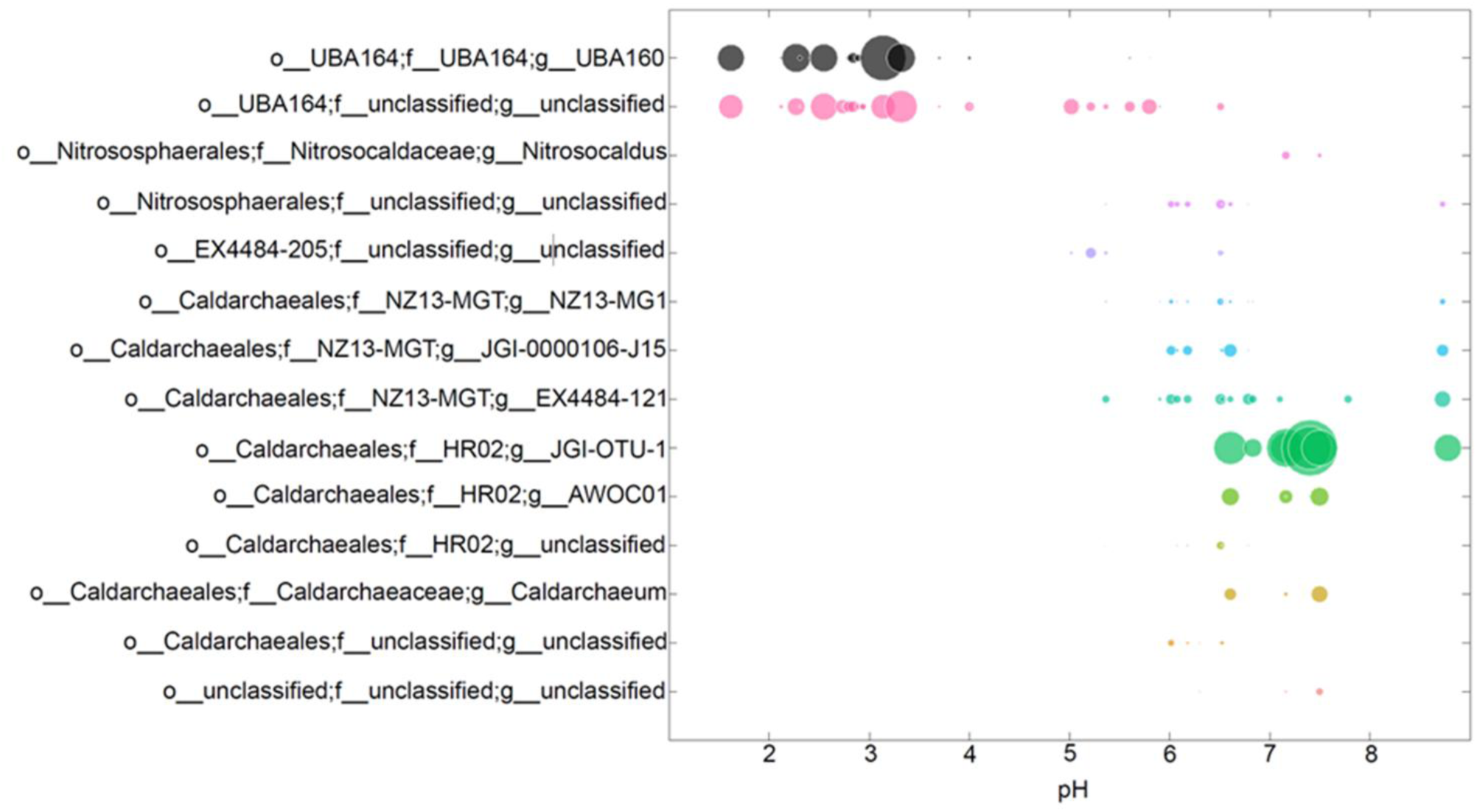
Relative abundances of *Nitrososphaeria* groups in Yellowstone thermal springs (data from Colman et al., 2024). Data are from 1022 metagenome-assembled genomes (MAGs) of 34 high-temperature, chemosynthetic springs in Yellowstone National Park and 444 MAGs from 35 published metagenomes. Nineteen of the sites from the current study overlap with those in Colman et al., 2024.

**Figure S3.**
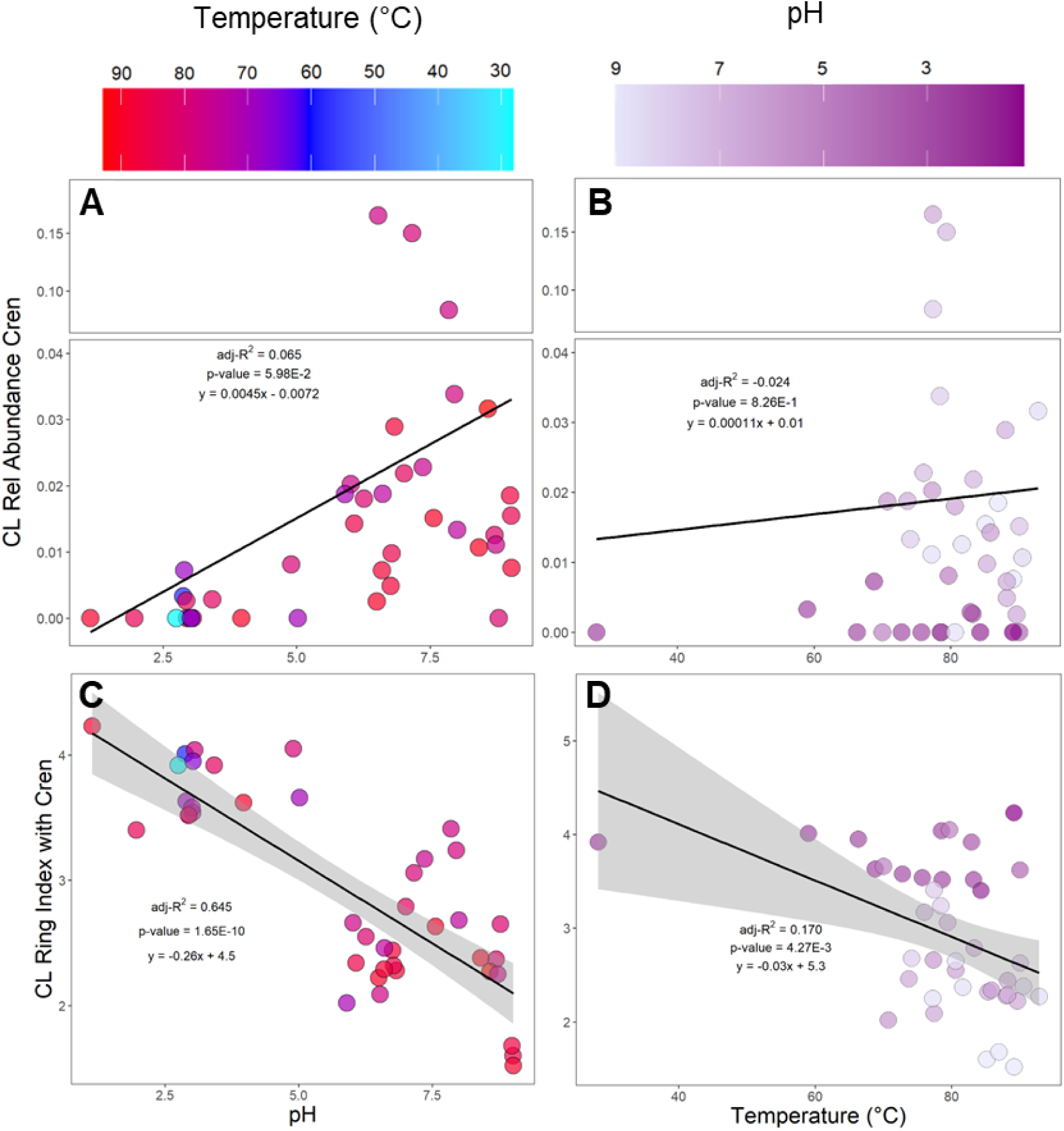
Relative abundances of CL crenarchaeol versus (A) pH and (B) temperature and Ring Index (with cren) versus (C) pH and (D) temperature for Yellowstone samples (N = 41) with fill by the other variable. Equations of lines of best fit, p-values, and adjusted-R^2^ values are included. A y-axis break and scale change are utilized to include high abundance outliers of crenarchaeol abundance while visualizing detailed distributions of the majority of samples. Significant RI linear relationships have 95% confidence intervals shaded in gray.

**Figure S4.**
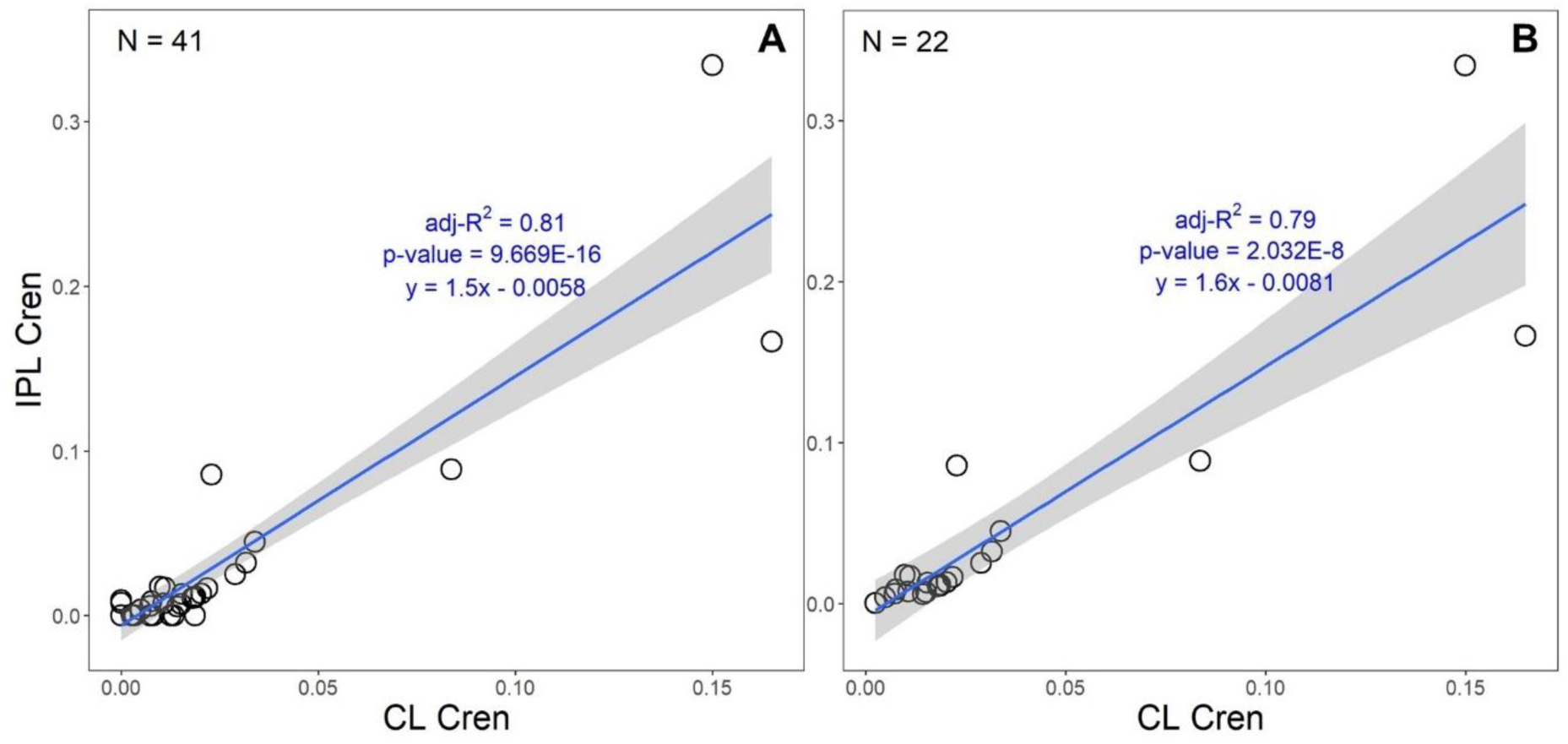
Linear regression model of IPL versus CL crenarchaeol relative abundance for each Yellowstone site. Panel (A) includes all sites while panel (B) excludes sites with zero values for CL or IPL crenarchaeol. The 95% confidence interval is shaded in gray and summary statistics are included.

**Figure S5.**
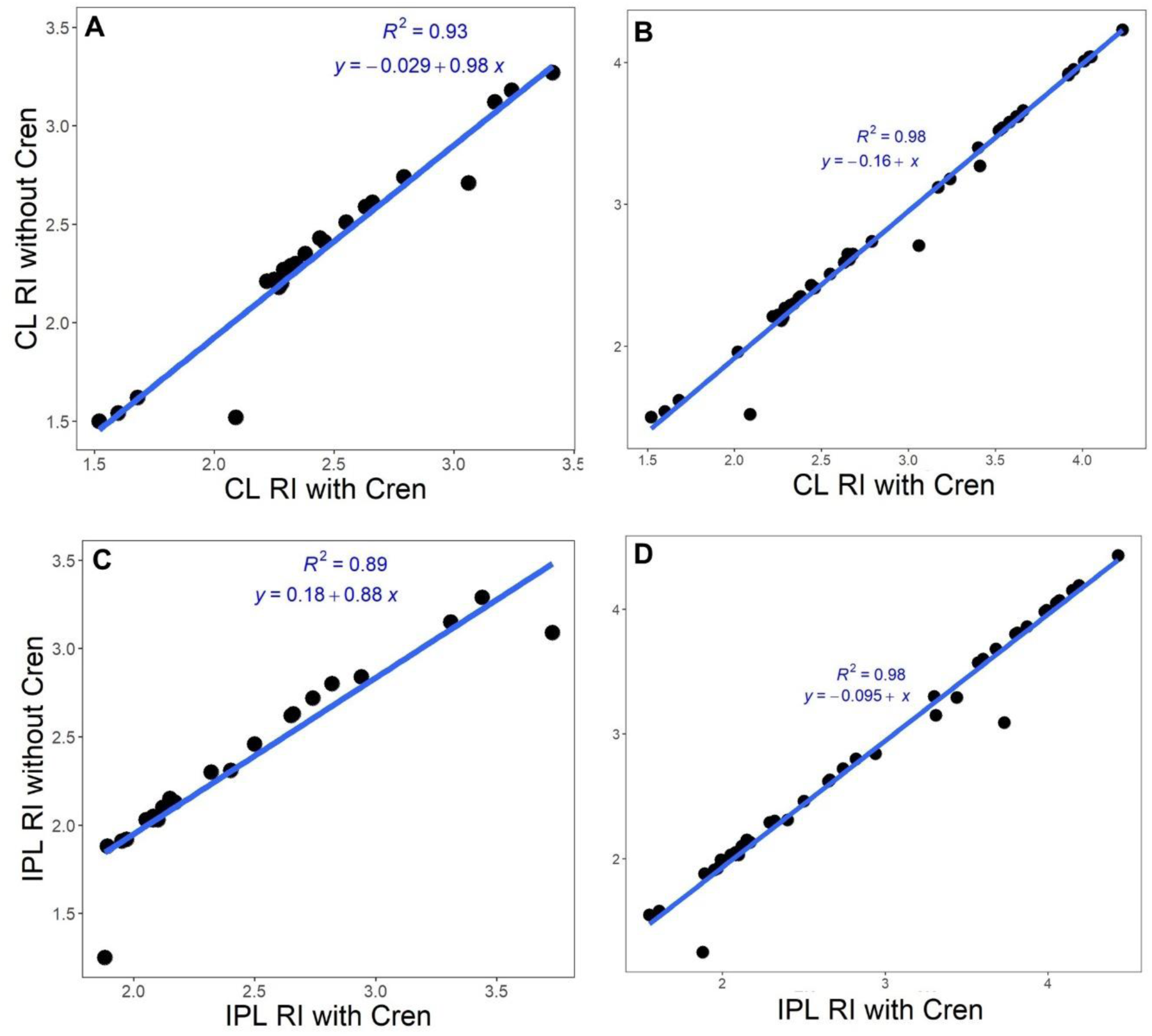
Simple linear regressions between Ring Index calculated without and with crenarchaeol for CL and IPL GDGT fractions from Yellowstone samples (See Equations 1 and 2). Panels (A) and (B) represent thermal springs with detectable amounts of crenarchaeol (N = 30), while Panels (C) and (D) have values from all 41 thermal spring samples regardless of crenarchaeol presence.

**Figure S6.**
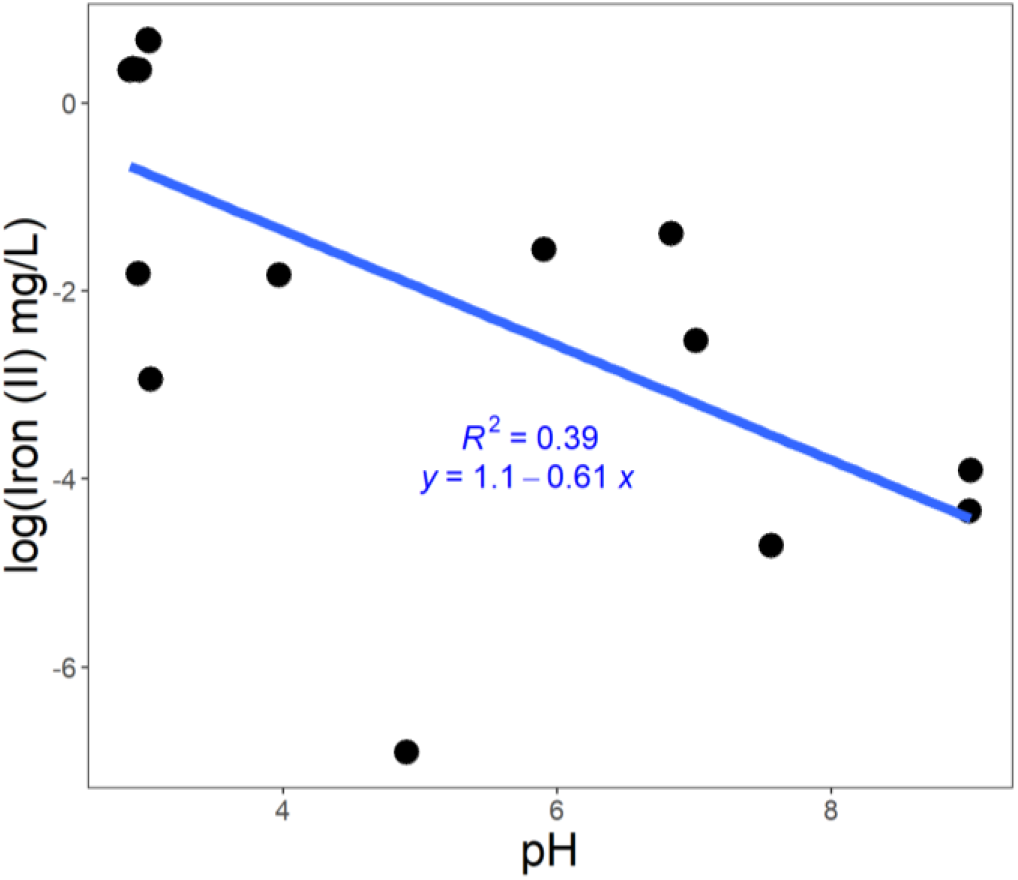
Simple linear regression of the log of Fe (II) concentration with pH for thermal springs with data for both parameters (N = 31). Non-finite log values are excluded, and the Spearman’s rho correlation p-value is 9.73 x 10^-5^.

## SUPPLEMENTAL TABLES

### Yellowstone Crenarchaeol

**Table S1.** Field geochemical and geophysical parameters, crenarchaeol relative abundances, and ring indices for 41 samples from Yellowstone National Park collected during field seasons from 2018-2022.

| Site Name | Year Sampled | Latitude (decimal degrees) | Longitude (decimal degrees) | pH | Temperature (°C) | Specific Conductivity (µS/cm) | Oxidation Reduction Potential (mV) | Dissolved Oxygen (mg/L) | Fe<sup>2+</sup> (mg/L) | S<sup>2-</sup> (µg/L) | CL Cren | CL Cren' | CL Cren+ Cren' | CL RI With Cren | CL RI Without Cren | IPL Cren | IPL Cren' | IPL Cren+ Cren' | IPL RI With Cren | IPL RI Without Cren |
| --- | --- | --- | --- | --- | --- | --- | --- | --- | --- | --- | --- | --- | --- | --- | --- | --- | --- | --- | --- | --- |
| Perpetual Spouter | 2018 | 44.7265959 | -110.709133 | 7.0 | 83 | 2790 | -120 | 0.8 | 0.08 | 10 | 0.019 | 0.0025 | 0.022 | 2.79 | 2.74 | 0.015 | 0.0017 | 0.016 | 2.50 | 2.46 |
| NSHPNN035 Main | 2018 | 44.7314663 | -110.711362 | 3.0 | 79 | 2120 | -100 | 0.1 | 1.42 | 1580 | BDL | BDL | BDL | 3.52 | 3.52 | BDL | BDL | BDL | 3.99 | 3.99 |
| NSHPNN035 Upper | 2018 | 44.7316004 | -110.711316 | 2.9 | 69 | 2130 | -40 | 0.7 | 1.44 | 470 | 0.0072 | BDL | 0.0072 | 3.63 | 3.62 | BDL | BDL | BDL | 3.99 | 3.99 |
| NSHPNN035 | 2018 | 44.7315187 | -110.711363 | 2.9 | 59 | 2170 | 210 | 2.0 | 1.42 | 170 | 0.0033 | BDL | 0.0033 | 4.01 | 4.01 | BDL | BDL | BDL | 3.30 | 3.30 |
| NHSPNN042 1 | 2018 | 44.7318275 | -110.711154 | 3.0 | 76 | 2040 | -90 | 0.2 | 1.94 | 1080 | BDL | BDL | BDL | 3.54 | 3.54 | BDL | BDL | BDL | 3.80 | 3.80 |
| NHSPNN042 2 | 2018 | 44.7318232 | -110.71092 | 3.0 | 73 | 2020 | -90 | 0.6 | 1.97 | 600 | BDL | BDL | BDL | 3.58 | 3.58 | BDL | BDL | BDL | 3.57 | 3.57 |
| NHSPNN042 3 | 2018 | 44.7317765 | -110.710901 | 3.1 | 79 | 1930 | -130 | 0.1 | NA | NA | BDL | BDL | BDL | 4.04 | 4.04 | BDL | BDL | BDL | 4.05 | 4.05 |
| Unnamed Norris Geyser Basin Spring | 2018 | 44.7318089 | -110.711018 | 2.8 | 28 | 2490 | 160 | 0.5 | NA | NA | BDL | BDL | BDL | 3.92 | 3.92 | 0.0026 | 0.0052 | 0.0078 | 3.87 | 3.86 |
| Cinder Pool (East Edge) | 2018 | 44.7324835 | -110.709773 | 4.0 | 90 | 2500 | -220 | 0.2 | 0.16 | 430 | BDL | BDL | BDL | 3.62 | 3.62 | BDL | BDL | BDL | 3.98 | 3.98 |
| Unnamed Midway Geyser Basin Spring 1 | 2018 | 44.5211885 | -110.811781 | 9.0 | 85 | 2040 | -320 | 0.5 | 0.01 | 10 | 0.0078 | 0.0077 | 0.015 | 1.60 | 1.54 | 0.0037 | 0.0093 | 0.013 | 2.17 | 2.13 |
| Unnamed Midway Geyser Basin Spring 2 | 2018 | 44.5198758 | -110.810763 | 9.0 | 89 | 2120 | -330 | 0.6 | 0.02 | 1290 | 0.0052 | 0.0024 | 0.0076 | 1.52 | 1.50 | 0.0023 | 0.0062 | 0.0086 | 2.32 | 2.30 |
| Unnamed Midway Geyser Basin Spring 3 | 2018 | 44.519861 | -110.810686 | 4.9 | 80 | 140 | -60 | 1.1 | 0.00 | 0 | 0.0081 | BDL | 0.0081 | 4.05 | 4.04 | BDL | BDL | BDL | 3.81 | 3.81 |
| Scaffolding Pool | 2018 | 44.5190822 | -110.809149 | 8.7 | 82 | 1760 | -130 | 0.8 | 0.00 | 0 | 0.011 | 0.0011 | 0.013 | 2.37 | 2.34 | BDL | BDL | BDL | 1.99 | 1.99 |
| Unnamed Midway Geyser Basin Spring 4 | 2018 | 44.51719 | -110.822738 | 7.6 | 90 | 1850 | -270 | 0.4 | 0.01 | 40 | 0.013 | 0.0017 | 0.015 | 2.63 | 2.59 | 0.0072 | BDL | 0.0072 | 2.82 | 2.80 |
| Beryl Spring | 2019 | 44.6787113 | -110.746493 | 6.8 | 88 | 2510 | -260 | 0.5 | 0.00 | 153 | 0.0049 | BDL | 0.0049 | 2.44 | 2.43 | 0.0026 | 0.0011 | 0.0036 | 1.89 | 1.88 |
| Bijah Spring | 2019 | 44.7611 | -110.73097 | 7.4 | 76 | 1470 | -30 | 1.6 | 0.00 | 27 | 0.021 | 0.0022 | 0.023 | 3.17 | 3.12 | 0.078 | 0.0080 | 0.086 | 3.44 | 3.29 |
| NEPSNN6 | 2019 | 44.72041 | -110.71326 | 2.9 | 83 | 1180 | 270 | 0.3 | 0.16 | 17 | 0.0026 | BDL | 0.0026 | 3.52 | 3.52 | BDL | BDL | BDL | 3.68 | 3.68 |
| GGCGNN017 | 2019 | 44.68406 | -110.74355 | 6.8 | 88 | 50 | -240 | 0.5 | 0.25 | 87 | 0.024 | 0.0053 | 0.029 | 2.28 | 2.20 | 0.014 | 0.011 | 0.025 | 2.10 | 2.03 |
| Unnamed Gibbon Canyon Spring | 2019 | 44.73616 | -110.70644 | 1.1 | 89 | 4470 | 310 | 0.2 | NA | 213 | BDL | BDL | BDL | 4.23 | 4.23 | BDL | BDL | BDL | 4.43 | 4.43 |
| Driveway Spring | 2022 | 44.4626324 | -110.852994 | 8.0 | 78 | 2200 | -100 | 1.1 | 0.00 | 0 | 0.034 | BDL | 0.034 | 3.24 | 3.18 | 0.041 | 0.0043 | 0.045 | 2.94 | 2.84 |
| Pentagonal Spring | 2022 | 44.463439 | -110.852837 | 7.9 | 77 | 2230 | -120 | 1.9 | 0.00 | 0 | 0.068 | 0.016 | 0.084 | 3.41 | 3.27 | 0.089 | BDL | 0.089 | 3.31 | 3.15 |
| Beryl Runoff | 2022 | 44.6786827 | -110.746465 | 6.8 | 85 | 2520 | -290 | 0.7 | NA | NA | 0.0058 | 0.0041 | 0.010 | 2.32 | 2.29 | 0.0070 | 0.011 | 0.018 | 1.97 | 1.92 |
| Beryl Main Overhang | 2022 | 44.6786622 | -110.746512 | 6.5 | 90 | 2510 | -330 | 0.4 | NA | NA | 0.0025 | BDL | 0.0025 | 2.22 | 2.21 | 0.0003 | BDL | 0.0003 | 2.15 | 2.15 |
| Beryl Side | 2022 | 44.6787151 | -110.746526 | 6.6 | 88 | 2520 | -310 | 0.5 | NA | NA | 0.0051 | 0.0022 | 0.0072 | 2.29 | 2.27 | 0.0009 | 0.0050 | 0.0059 | 2.12 | 2.10 |
| GGSN067 | 2020 | 44.69032 | -110.72993 | 2.0 | 84 | 3410 | 330 | 0.4 | NA | 47 | BDL | BDL | BDL | 3.40 | 3.40 | BDL | BDL | BDL | 4.19 | 4.19 |
| Unnamed Geyser Creek Spring 1 | 2020 | 44.69067 | -110.72977 | 6.1 | 86 | 5380 | 0 | 0.0 | BDL | 515 | 0.0087 | 0.0056 | 0.014 | 2.34 | 2.30 | 0.0016 | 0.0040 | 0.0056 | 2.05 | 2.03 |
| Unnamed Geyser Creek Spring 2 | 2020 | 44.69038 | -110.7289 | 6.0 | 77 | 5090 | 50 | 0.2 | BDL | 73 | 0.017 | 0.0035 | 0.020 | 2.66 | 2.61 | 0.0094 | 0.0038 | 0.013 | 1.95 | 1.91 |
| SMHNN007 | 2020 | 44.7536 | -110.40418 | 6.5 | 77 | 2000 | 30 | 0.1 | NA | 2500 | 0.13 | 0.034 | 0.17 | 2.09 | 1.52 | 0.12 | 0.049 | 0.17 | 1.88 | 1.25 |
| Bone Pool | 2020 | 44.68915 | -110.7285 | 6.6 | 74 | 5180 | 20 | 0.8 | BDL | 16 | 0.013 | 0.0057 | 0.019 | 2.46 | 2.41 | 0.0087 | 0.0027 | 0.011 | 2.65 | 2.62 |
| GGSN064 | 2020 | 44.69033 | -110.72935 | 3.4 | 83 | 2470 | 240 | 0.2 | NA | 150 | 0.0029 | BDL | 0.0029 | 3.92 | 3.91 | BDL | BDL | BDL | 4.15 | 4.15 |
| Unnamed Solfatara Plateau Spring | 2020 | 44.75154 | -110.4151 | 8.0 | 74 | 4110 | -70 | 1.0 | BDL | 2570 | 0.012 | 0.0014 | 0.013 | 2.68 | 2.65 | BDL | BDL | BDL | 1.55 | 1.55 |
| Obsidian Pool | 2019 | 44.61006 | -110.43882 | 5.9 | 71 | 1120 | 90 | NA | 0.21 | 130 | 0.0069 | 0.012 | 0.019 | 2.02 | 1.96 | BDL | BDL | BDL | 3.60 | 3.60 |
| NHSPNN155 | 2019 | 44.73616 | -110.70649 | 5.0 | 70 | 3640 | NA | 0.0 | NA | 7858 | BDL | BDL | BDL | 3.66 | 3.66 | 0.0019 | 0.0072 | 0.0092 | 1.61 | 1.58 |
| Boulder Pool | 2019 | 44.55878 | -110.844 | 8.4 | 91 | 4040 | -100 | 0.0 | BDL | 2190 | 0.0030 | 0.0077 | 0.011 | 2.38 | 2.35 | 0.0018 | 0.0055 | 0.0073 | 2.74 | 2.72 |
| LRNN012 | 2019 | 44.56153 | -110.83685 | 6.3 | 81 | 3700 | 50 | 0.0 | BDL | 319 | 0.0078 | 0.010 | 0.018 | 2.55 | 2.51 | 0.0036 | 0.0070 | 0.011 | 2.66 | 2.63 |
| MNN017 | 2019 | 44.51987 | -110.81077 | 9.0 | 87 | 4370 | -130 | 0.3 | BDL | 1190 | 0.0096 | 0.0089 | 0.019 | 1.68 | 1.62 | 0.0039 | 0.0071 | 0.011 | 2.08 | 2.05 |
| Mound Spring | 2019 | 44.56485 | -110.86005 | 8.6 | 93 | 3830 | -110 | 0.0 | BDL | 1092 | 0.0095 | 0.022 | 0.032 | 2.27 | 2.18 | BDL | 0.032 | 0.032 | 2.40 | 2.31 |
| MVN002 | 2019 | 44.60993 | -110.43939 | 3.0 | 66 | 4120 | 260 | NA | 0.05 | 13 | BDL | BDL | BDL | 3.95 | 3.95 | BDL | BDL | BDL | 4.07 | 4.07 |
| Scaffolding Pool | 2019 | 44.51914 | -110.80914 | 8.8 | 81 | 3600 | -120 | 1.2 | BDL | 23 | BDL | BDL | BDL | 2.65 | 2.65 | BDL | BDL | BDL | 2.29 | 2.29 |
| MRCHSGNN093 | 2019 | 44.50889 | -110.80895 | 7.16 | 79 | 910 | 0 | 1.0 | BDL | BDL | 0.15 | BDL | 0.15 | 3.06 | 2.71 | 0.30 | 0.036 | 0.33 | 3.73 | 3.09 |
| Azure Pool | 2019 | 44.56097 | -110.83286 | 8.73 | 77 | 3450 | -120 | 0.8 | BDL | 345 | 0.0018 | 0.0093 | 0.011 | 2.25 | 2.22 | BDL | 0.017 | 0.017 | 2.08 | 2.03 |

**Table S2.**
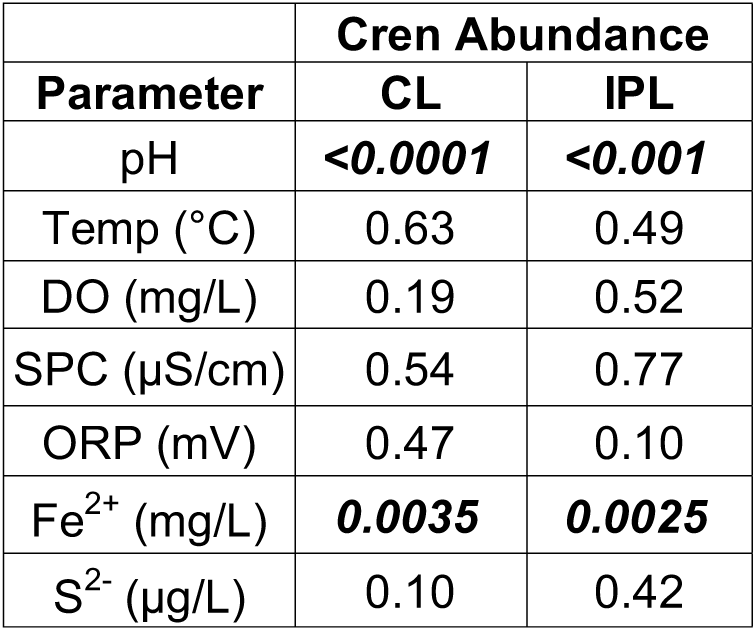
Spearman’s rho correlation p-values for environmental parameters and crenarchaeol relative abundance from Yellowstone samples. Analyses are run for both core (CL) and intact polar lipid (IPL) fractions and significant correlations are indicated in bold italics.

| Parameter | Cren Abundance |  |
| --- | --- | --- |
|  | CL | IPL |
| pH | **<0.0001** | **<0.001** |
| Temp (°C) | 0.63 | 0.49 |
| DO (mg/L) | 0.19 | 0.52 |
| SPC (μS/cm) | 0.54 | 0.77 |
| ORP (mV) | 0.47 | 0.10 |
| Fe<sup>2+</sup> (mg/L) | **0.0035** | **0.0025** |
| S<sup>2-</sup> (μg/L) | 0.10 | 0.42 |

**Table S3:** Outputs of simple linear regression models for environmental parameters and CL and IPL relative abundance of crenarchaeol from Yellowstone samples. The significance threshold is set to a p-value of 0.05 and any significant correlations would be indicated in bold italics.

| Parameter | Crenarchaeol Abundance |  |  |  |
| --- | --- | --- | --- | --- |
|  | CL |  | IPL |  |
|  | p-value | adjusted R<sup>2</sup> | p-value | adjusted R<sup>2</sup> |
| pH | 0.06 | 0.07 | 0.15 | 0.03 |
| Temp (°C) | 0.83 | -0.02 | 0.98 | -0.03 |
| DO (mg/L) | 0.34 | 0.00 | 0.17 | 0.03 |
| SPC (μS/cm) | 0.18 | 0.02 | 0.09 | 0.05 |
| ORP (mV) | 0.99 | -0.03 | 0.91 | -0.03 |
| Fe<sup>2+</sup> (mg/L) | 0.14 | 0.04 | 0.32 | 0.00 |
| S<sup>2-</sup> (μg/L) | 0.91 | -0.03 | 0.92 | -0.03 |

**Table S4.** Outputs of 10 multiple linear regression models with highest adjusted R^2^ values for environmental parameters and relative abundances of CL crenarchaeol from Yellowstone samples. The significance threshold is set to 0.05 and significant p-values are indicated in bold italics. Models are reported in order of decreasing adjusted R^2^ values.

| Input Parameters | Crenarchaeol Abundance |  |  |
| --- | --- | --- | --- |
| | p-value | Multiple $R^2$ | Adjusted $R^2$ |
| pH, SPC, ORP, $S^{2-}$ | ***2.33E-02*** | 0.31 | 0.21 |
| pH, SPC, ORP, DO, $S^{2-}$ | 5.12E-02 | 0.32 | 0.20 |
| pH, Temp, SPC, ORP, $S^{2-}$ | ***4.80E-02*** | 0.31 | 0.19 |
| pH, SPC, ORP | ***1.47E-02*** | 0.25 | 0.19 |
| pH, SPC, ORP, DO | ***2.77E-02*** | 0.27 | 0.19 |
| pH, Temp, SPC, ORP, DO, $S^{2-}$ | 7.66E-02 | 0.34 | 0.18 |
| pH, Temp, SPC, ORP | ***3.46E-02*** | 0.25 | 0.17 |
| pH, Temp, SPC, ORP, DO | 5.55E-02 | 0.28 | 0.16 |
| pH, SPC, ORP, $Fe^{2+}$ | 1.17E-01 | 0.24 | 0.12 |
| DO, $Fe^{2+}$ | 8.61E-02 | 0.17 | 0.11 |

### Yellowstone Ring Index

**Table S5.** Spearman’s rho correlation p-values for environmental parameters and Ring Index from Yellowstone samples. Analyses are run for both core (CL) and intact polar lipid (IPL) fractions and significant correlations are indicated in bold italics.

| Parameter | Ring Index |  |
| --- | --- | --- |
|  | CL | IPL |
| pH | ***<0.0001*** | ***<0.0001*** |
| Temp ($^{\circ}C$) | ***0.0060*** | 0.34 |
| DO (mg/L) | 0.67 | 0.93 |
| SPC ($\mu S/cm$) | 0.64 | 0.10 |
| ORP (mV) | ***<0.001*** | ***0.0019*** |
| $Fe^{2+}$ (mg/L) | ***0.018*** | ***<0.001*** |
| $S^{2-}$ ($\mu g/L$) | 0.43 | 0.16 |

**Table S6.** Outputs of simple linear regression models for environmental parameters and CL and IPL Ring Index with crenarchaeol from Yellowstone samples. The significance threshold is set to a p-value of 0.05 and significant correlations are indicated in bold italics.

| Parameter | Ring Index |  |  |  |
| --- | --- | --- | --- | --- |
|  | CL |  | IPL |  |
|  | p-value | adjusted R<sup>2</sup> | p-value | adjusted R<sup>2</sup> |
| pH | **<0.0001** | 0.64 | **<0.0001** | 0.55 |
| Temp (°C) | **0.0043** | 0.17 | 0.08 | 0.05 |
| DO (mg/L) | 0.38 | -0.01 | 0.76 | -0.02 |
| SPC (μS/cm) | 0.40 | -0.01 | 0.08 | 0.05 |
| ORP (mV) | **<0.001** | 0.31 | **<0.001** | 0.31 |
| Fe<sup>2+</sup> (mg/L) | **0.0042** | 0.22 | **0.0046** | 0.22 |
| S<sup>2-</sup> (μg/L) | 0.97 | -0.03 | **0.028** | 0.11 |

**Table S7.** Outputs of 10 multiple linear regression models with highest adjusted R^2^ values for environmental parameters and CL Ring Indices with crenarchaeol from Yellowstone samples. The significance threshold is set to 0.05 and significant p-values are indicated in bold italics. Models are listed in order of decreasing adjusted R^2^ values.

| Input Parameters | Ring Index |  |  |
| --- | --- | --- | --- |
|  | p-value | Multiple R<sup>2</sup> | Adjusted R<sup>2</sup> |
| pH, Temp, DO, Fe<sup>2+</sup> | **<0.0001** | 0.82 | 0.80 |
| pH, DO, Fe<sup>2+</sup> | **<0.0001** | 0.82 | 0.79 |
| pH, Temp, DO, Fe<sup>2+</sup>, S<sup>2-</sup> | **<0.0001** | 0.83 | 0.79 |
| pH, SPC, DO, Fe<sup>2+</sup> | **<0.0001** | 0.82 | 0.79 |
| pH, Temp, ORP, DO, Fe<sup>2+</sup> | **<0.0001** | 0.83 | 0.79 |
| pH, SPC, DO, Fe<sup>2+</sup>, S<sup>2-</sup> | **<0.0001** | 0.83 | 0.79 |
| pH, DO, Fe<sup>2+</sup>, S<sup>2-</sup> | **<0.0001** | 0.82 | 0.79 |
| pH, Temp, ORP, DO, S<sup>2-</sup> | **<0.0001** | 0.82 | 0.79 |
| pH, Temp, DO, S<sup>2-</sup> | **<0.0001** | 0.81 | 0.79 |
| pH, Temp, SPC, DO, Fe<sup>2+</sup> | **<0.0001** | 0.82 | 0.79 |

## Supplementary Information

### Comparisons#of Core and Polar Fractions

To identify whether studies should split GDGTs into CL and IPL fractions, we compared our YNP results for these two fractions. CL and IPL crenarchaeol relative abundances are strongly positively correlated (R^2^ = 0.81; Figure S4). Interestingly, the slope of this relationship is 1.5, indicating that on average, crenarchaeol has greater relative abundance in IPL fractions than in CL fractions. IPL lipids are thought to derive from extant microbial communities and CL lipids record long-term production, indicating that microbial communities in sampled YNP springs may have been producing more crenarchaeol upon collection than the time integral represented in the CL fraction (Lipp and Hinrichs, 2009). Another hypothesis is that crenarchaeol is more labile than the other GDGTs and its CL abundance decreases over time relative to other GDGTs. To distinguish between these two hypotheses, we plotted IPL Ring Index with crenarchaeol versus CL Ring Index with crenarchaeol, which produced a linear trendline with the equation y = 0.88x + 0.35 (R^2^ = 0.58). The result of plotting IPL RI without cren versus CL RI without cren is almost identical with a trendline of y = 0.88x + 0.34 (R^2^ = 0.60). The slopes of 0.88 are close to a 1:1 relationship, different from the crenarchaeol abundance slope of 1.5 for IPL versus CL. This indicates that crenarchaeol may be preferentially degraded compared to other GDGTs with just cyclopentyl rings, supporting the degradation hypothesis over the environmental change hypothesis.

Four sites deviate from the best-fit line and lie outside of the 95% confidence interval in Figure S4, but these outliers maintain a positive relationship between CL and IPL crenarchaeol. The four samples all have relatively high CL and IPL crenarchaeol relative abundances for the Yellowstone dataset. Two sites, SMHNN007 (CL = 0.17; IPL = 0.17) and Pentagonal Spring (CL = 0.08; IPL = 0.09) have more CL crenarchaeol than predicted by the best-fit line with a slope of 1.5. Both springs have similar relative abundances of crenarchaeol in CL and IPL fractions, indicating the native microbial cells were producing their time-averaged proportion of crenarchaeol at the time of sampling from potentially stable geochemical conditions. Another possibility is that IPL crenarchaeol degradation is enhanced in these springs.

The other two high relative abundance sites are MRCHSGNN093 (CL = 0.15; IPL = 0.33) and Bijah Spring (CL = 0.02; IPL = 0.09), which have more IPL crenarchaeol than predicted by the best fit line. The greater relative abundance of IPL crenarchaeol indicates that either the degradation rate of IPL crenarchaeol is lower in these springs, or that the microbial community at the time of sampling of these springs was producing more crenarchaeol relative to other GDGTs than the historical average of this spring. This phenomenon may indicate geochemical conditions that inhibit IPL crenarchaeol degradation, seasonal variation in environmental parameters that influence crenarchaeol relative abundance, or recent shifts in microbial community or geochemical/geophysical conditions of the springs. Hydrothermal conditions of springs are known to vary with major geyser eruptions, hydrologic changes, earthquakes, discharge rates, variable mixing of gases/waters, and annual and seasonal disturbances, which could explain these differences in CL and IPL crenarchaeol (Heasler and Jaworowski, 2018).

Conditions experienced by microorganisms in thermal spring environments are also known to be influenced by the volcanic system that drives hydrothermal activity in Yellowstone National Park, which undergoes complex fluctuations over time (Hurwitz and Lowenstern, 2014). Climate variation may also play a role in altering thermal spring temperature and chemistry, which would influence trends observed for the 41 Yellowstone samples collected during different times of the year. Variations over even smaller temporal scales can influence thermal spring conditions. Samples were collected from thermal springs across Norris Geyser Basin, which is known to have ground temperature variation from ∼30°C to ∼50°C over the course of a single day (Neale et al., 2016). Long-term increases in annual mean air temperature may represent more systemic changes for spring chemistry through indirect processes such as precipitation and nutrient flux, while drought conditions alter input from water sources such as volcanically heated reservoirs, riverine input, and precipitation (Heeter et al., 2021). During the 2022 field season, sediment samples were collected shortly after significant snowfall, which may have influenced spring geochemistry both by direct input and alteration of water table level. Differences in source-water depths of these hydrothermal systems also have important implications for the geochemistry of these springs (Fournier et al., 2002).

The variable conditions of thermal springs in Yellowstone may explain discrepancies between crenarchaeol presence/absence in CL and IPL fractions from the same sample. Out of 41 samples generated in this study, eight have detectable levels of CL crenarchaeol and no detectable levels of IPL crenarchaeol (see Excel file of data table). Interestingly, two have detectable levels of IPL crenarchaeol and no detectable levels of CL crenarchaeol. Samples with measurable amounts of CL crenarchaeol and non-detectable IPL crenarchaeol are expected and can be explained by the cessation of crenarchaeol production in these springs with environmental variation or microbial community change because core lipids are degradation products of intact polar lipids. In contrast, the presence of detectable amounts of IPL crenarchaeol and no CL crenarchaeol in two sites is more difficult to explain. One of these sites is a temperature outlier, Unnamed Norris Geyser Basin Spring (28.2°C, pH 2.75), which may have recently begun crenarchaeol production and has a slow degradation rate of archaeal lipids at the relatively low temperature of this spring. The other anomaly, NHSPNN155 (70°C, pH 5.02), does not support this slow-degradation interpretation. In NHSPNN155, the only anomaly is the high dissolved sulfide level, at 2.45 x 10^-4^ M. This is the highest value of the Yellowstone dataset, with a second-highest concentration being 8.01 x 10^-5^ M. A majority (75%) of samples with S^2-^ data available (36 sites) are at least an order of magnitude lower with an average concentration of 700 µg L^-1^. Upon removal of NHSPNN155 from this dataset average, the value drops to 496 µg L^-1^. Sulfide concentration was not measured for Unnamed Norris Geyser Basin Spring, which prevents comparison of sulfur abundance across these two sites. However, lipid sulfurization is a known diagenetic process that may lead to enhanced degradation of CL GDGTs in the presence of high sulfur concentrations, which could lead to non-detectable levels of CL crenarchaeol (Schouten et al., 2013; Adam et al., 2000). While there are exceptions, the majority of samples (75.6%) have consistent crenarchaeol presence and absence across CL and IPL fractions.

The strong linear relationship between CL and IPL crenarchaeol relative abundances justifies plotting results of only CL fraction analyses because similar statistical results are observed for IPL fractions. Multiple linear regression analyses were also run for IPL crenarchaeol to confirm similarity of results with those for CL crenarchaeol reported in Table S3. While results should do not vary between fractions, we chose to plot results for CL crenarchaeol rather than IPL crenarchaeol because CL lipids record a longer history of GDGT production that is not as subject to recent seasonal changes as expected for IPL crenarchaeol (Lipp and Hinrichs, 2009). In addition, we utilized internal GTGT standard to calculate rough absolute abundances of GDGTs in the samples. CL fractions represent the majority of the GDGTs from 94% of samples with internal standard (N = 33), making them preferable for robust quantification on the QQQ and use in analyses. Two samples, Unnamed Midway Geyser Basin Spring 2 and Driveway Spring had more IPL than CL GDGTs (57% and 84%, respectively), which is anomalous in this dataset.

The other 31 samples whose absolute abundance could be calculated had an average percent CL composition of 85.9% ± 13.0, establishing a significant dominance of CL lipids in these samples. The rough calculations from addition of GTGT standard are imprecise and are best suited for order-of-magnitude calculations, which prevents accurate combinations of absolute GDGT abundances from IPL and CL data. Past studies have varied in the lumping or splitting of CL and IPL fractions, so we report compiled literature samples as either total isoprenoid-GDGT abundances or CL abundances (Boyd et al., 2013; Xie et al., 2015). In comparing the literature values to the new data generated in this study, we present only CL crenarchaeol data due to the higher magnitude of CL versus IPL lipid absolute abundances and representation of a long record of GDGT production.

While differences between CL and IPL crenarchaeol relative abundances can indicate recent shifts in microbial community or environmental conditions in each spring, broad trends of cyclopentyl ring abundances (represented by RI) are consistent across CL and IPL fractions (see Table S4 and Table S5). Given this observation, it may be sufficient to analyze relative GDGT abundances amongst total isoprenoid-GDGTs, rather than splitting into CL and IPL fractions, depending on the scientific goal. This is significant because splitting samples and processing fractions in tandem involves labor intensive wet chemistry protocols and doubles analytical machine time. However, if the environment in question has undergone significant recent changes or researchers are interested in a lipid of high abundance, fraction splitting is suggested to obtain a comprehensive view of lipid production.

### Effect of Crenarchaeol on Ring Index Calculations

We compared YNP Ring Index calculated with and without crenarchaeol in Supplemental Figure S5 and demonstrate a strong 1:1 correlation between the two, justifying our utilization of RI calculated with crenarchaeol subsequent analyses. Panels A and C show regressions of YNP samples with detectable crenarchaeol for CL (R^2^ = 0.93) and IPL fractions (R^2^ = 0.89), indicating a strong correlation of RI values. The slopes of the significant CL and IPL correlations are 0.98 and 0.88, respectively, which are close to the 1:1 relationship expected for RI values from the same sample. Two samples lie outside of the 95% confidence interval of the regression relationships, which are from MRCHSGNN093 and SMHNN007, the springs with the highest relative abundances of both CL and IPL crenarchaeol in the Yellowstone dataset (CL = 0.15; IPL = 0.33 and CL = 0.17; IPL = 0.17, respectively). As expected, higher relative abundances of crenarchaeol increase discrepancies between the two RIs since crenarchaeol contributes four (cyclopentyl) rings to the RI with crenarchaeol. Adding crenarchaeol to the RI of MRCHSGNN093 increased CL RI from 2.71 to 3.06 and IPL from 3.09 to 3.73, while the RI of SMHNN007 increased from 1.52 to 2.09 for CL and 1.25 to 1.88 for IPL. These results indicate that incorporating crenarchaeol into RI calculations must be thought through for springs with high concentrations of crenarchaeol.

Depending on the intended purpose of Ring Index, crenarchaeol may best be excluded from RI calculations. In marine sediments, RI can be used to determine whether the TEX_86_ temperature estimates are influenced by factors other than temperature, because Ring Index and TEX_86_ values have a global correlation when temperature is the controlling factor (Zhang et al., 2016). However, if variables other than temperature are more important for ring cyclization, then the relationship between Ring Index and TEX_86_ temperatures fails. Temperature is associated with crenarchaeol distributions, so including crenarchaeol in RI for paleothermometry may be reasonable even if crenarchaeol increases membrane permeability if it can provide a nuanced perspective into GDGT cyclization responses to temperature. If RI is being used to indicate cyclopentyl-forming enzyme activity, crenarchaeol may be included as a four-ringed member. However, if one is using RI as a stress response or indictor of decreased permeability, it may be best to exclude crenarchaeol from ring index, because this lipid prefers moderate temperatures, supporting a membrane-expanding function.

## References

1. Abby, S. S., Kerou, M., & Schleper, C. (2020). Ancestral Reconstructions Decipher Major Adaptations of Ammonia-Oxidizing Archaea upon Radiation into Moderate Terrestrial and Marine Environments. mBio, 11(5), 10.1128/mbio.02371-20. 10.1128/mbio.02371-20

2. Adam, P., Schneckenburger, P., Schaeffer, P., & Albrecht, P. (2000). Clues to early diagenetic sulfurization processes from mild chemical cleavage of labile sulfur-rich geomacromolecules. Geochimica et Cosmochimica Acta, 64(20), 3485–3503. 10.1016/S0016-7037(00)00443-9

3. Albers, S. V., van de Vossenberg, J. L., Driessen, A. J., & Konings, W. N. (2000). Adaptations of the archaeal cell membrane to heat stress. Frontiers in Bioscience-Landmark, 5(3), 813–820.

4. Amenabar, M. J., & Boyd, E. S. (2018). Mechanisms of Mineral Substrate Acquisition in a Thermoacidophile. Applied and Environmental Microbiology, 84(12), e00334–18. 10.1128/AEM.00334-18f

5. Amend, J. P., & Shock, E. L. (2001). Energetics of overall metabolic reactions of thermophilic and hyperthermophilic Archaea and Bacteria. FEMS Microbiology Reviews, 25(2), 175–243. 10.1111/j.1574-6976.2001.tb00576.x

6. Beam, J. P., Jay, Z. J., Kozubal, M. A., & Inskeep, W. P. (2014). Niche specialization of novel Thaumarchaeota to oxic and hypoxic acidic geothermal springs of Yellowstone National Park. The ISME Journal, 8(4), 938–951. 10.1038/ismej.2013.193

7. Becker, K. W., Lipp, J. S., Zhu, C., Liu, X.-L., & Hinrichs, K.-U. (2013). An improved method for the analysis of archaeal and bacterial ether core lipids. Organic Geochemistry, 61, 34–44. 10.1016/j.orggeochem.2013.05.007

8. Bice, K. L., Birgel, D., Meyers, P. A., Dahl, K. A., Hinrichs, K.-U., & Norris, R. D. (2006). A multiple proxy and model study of Cretaceous upper ocean temperatures and atmospheric CO2 concentrations. Paleoceanography, 21(2). 10.1029/2005PA001203

9. Blewett, J., Naafs, B. D. A., Gallego-Sala, A. V., & Pancost, R. D. (2020). Effects of temperature and pH on archaeal membrane lipid distributions in freshwater wetlands. Organic Geochemistry, 148, 104080. 10.1016/j.orggeochem.2020.104080

10. Bligh, E. G., & Dyer, W. J. (1959). A rapid method of total lipid extraction and purification. Canadian Journal of Biochemistry and Physiology, 37(8), 911–917. 10.1139/o59-099

11. Blum, L. N., Colman, D. R., Eloe-Fadrosh, E. A., Kellom, M., Boyd, E. S., Zhaxybayeva, O., & Leavitt, W. D. (2022). Distribution and abundance of tetraether lipid cyclization genes in terrestrial thermal springs reflects pH (p. 2022.08.15.504015). bioRxiv. 10.1101/2022.08.15.504015

12. Boyd, E., Hamilton, T., Wang, J., He, L., & Zhang, C. (2013). The Role of Tetraether Lipid Composition in the Adaptation of Thermophilic Archaea to Acidity. Frontiers in Microbiology, 4. https://www.frontiersin.org/articles/10.3389/fmicb.2013.00062

13. Boyd, E. S., Pearson, A., Pi, Y., Li, W.-J., Zhang, Y. G., He, L., Zhang, C. L., & Geesey, G. G. (2011). Temperature and pH controls on glycerol dibiphytanyl glycerol tetraether lipid composition in the hyperthermophilic crenarchaeon Acidilobus sulfurireducens. Extremophiles, 15(1), 59–65. 10.1007/s00792-010-0339-y

14. Boyer, G. M., Schubotz, F., Summons, R. E., Woods, J., & Shock, E. L. (2020). Carbon Oxidation State in Microbial Polar Lipids Suggests Adaptation to thermal Spring Temperature and Redox Gradients. Frontiers in Microbiology, 11. https://www.frontiersin.org/articles/10.3389/fmicb.2020.00229

15. Brochier-Armanet, C., Boussau, B., Gribaldo, S., & Forterre, P. (2008). Mesophilic crenarchaeota: Proposal for a third archaeal phylum, the Thaumarchaeota. Nature Reviews Microbiology, 6(3), Article 3. 10.1038/nrmicro1852

16. Buessecker, S., Palmer, M., Lai, D., Dimapilis, J., Mayali, X., Mosier, D., Jiao, J.-Y., Colman, D. R., Keller, L. M., St. John, E., Miranda, M., Gonzalez, C., Gonzalez, L., Sam, C., Villa, C., Zhuo, M., Bodman, N., Robles, F., Boyd, E. S., … Dodsworth, J. A. (2022). An essential role for tungsten in the ecology and evolution of a previously uncultivated lineage of anaerobic, thermophilic Archaea. Nature Communications, 13(1), 3773. 10.1038/s41467-022-31452-8

17. Burgess, E. A., Unrine, J. M., Mills, G. L., Romanek, C. S., & Wiegel, J. (2012). Comparative Geochemical and Microbiological Characterization of Two Thermal Pools in the Uzon Caldera, Kamchatka, Russia. Microbial Ecology, 63(3), 471–489. 10.1007/s00248-011-9979-4

18. Church, M. J., Wai, B., Karl, D. M., & DeLong, E. F. (2010). Abundances of crenarchaeal amoA genes and transcripts in the Pacific Ocean. Environmental Microbiology, 12(3), 679–688. 10.1111/j.1462-2920.2009.02108.x

19. Clayton, T. D., & Byrne, R. H. (1993). Spectrophotometric seawater pH measurements: Total hydrogen ion concentration scale calibration of m-cresol purple and at-sea results. Deep Sea Research Part I: Oceanographic Research Papers, 40(10), 2115–2129. 10.1016/0967-0637(93)90048-8

20. Cobban, A., Zhang, Y., Zhou, A., Weber, Y., Elling, F. J., Pearson, A., & Leavitt, W. D. (2020). Multiple environmental parameters impact lipid cyclization in Sulfolobus acidocaldarius. Environmental Microbiology, 22(9), 4046–4056. 10.1111/1462-2920.15194

21. Colman, D. R., Feyhl-Buska, J., Robinson, K. J., Fecteau, K. M., Xu, H., Shock, E. L., & Boyd, E. S. (2016). Ecological differentiation in planktonic and sediment-associated chemotrophic microbial populations in Yellowstone thermal springs. FEMS Microbiology Ecology, 92(9), fiw137. 10.1093/femsec/fiw137

22. Colman, D. R., Keller, L. M., Arteaga-Pozo, E., Andrade-Barahona, E., St. Clair, B., Shoemaker, A., Cox, A., & Boyd, E. S. (2024). Covariation of thermal spring geochemistry with microbial genomic diversity, function, and evolution. Nature Communications, 15(1), 7506. 10.1038/s41467-024-51841-5

23. thermal Colman, D. R., Lindsay, M. R., Harnish, A., Bilbrey, E. M., Amenabar, M. J., Selensky, M. J., Fecteau, K. M., Debes II, R. V., Stott, M. B., Shock, E. L., & Boyd, E. S. (2021). Seasonal hydrologic and geologic forcing drive thermal spring geochemistry and microbial biodiversity. Environmental Microbiology, 23(7), 4034–4053. 10.1111/1462-2920.15617

24. Colman, D. R., Poudel, S., Hamilton, T. L., Havig, J. R., Selensky, M. J., Shock, E. L., & Boyd, E. S. (2018). Geobiological feedbacks and the evolution of thermoacidophiles. The ISME Journal, 12(1), 225–236. 10.1038/ismej.2017.162

25. Damer, B., & Deamer, D. (2020). The thermal Spring Hypothesis for an Origin of Life. Astrobiology, 20(4), 429–452. 10.1089/ast.2019.2045

26. Damsté, J. S. S., Schouten, S., Hopmans, E. C., Duin, A. C. T. van, & Geenevasen, J. A. J. (2002). Crenarchaeol. Journal of Lipid Research, 43(10), 1641–1651. 10.1194/jlr.M200148-JLR200

27. De La Torre, J. R., Walker, C. B., Ingalls, A. E., Könneke, M., & Stahl, D. A. (2008). Cultivation of a thermophilic ammonia oxidizing archaeon synthesizing crenarchaeol. Environmental Microbiology, 10(3), 810–818. 10.1111/j.1462-2920.2007.01506.x

28. De Rosa, M., & Gambacorta, A. (1988). The lipids of archaebacteria. Progress in Lipid Research, 27(3), 153–175. 10.1016/0163-7827(88)90011-2

29. De Rosa, M., Gambacorta, A., Nicolaus, B., Sodano, S., & Bu’Lock, J. D. (1980). Structural regularities in tetraether lipids of Caldariella and their biosynthetic and phyletic implications. Phytochemistry, 19(5), 833–836. 10.1016/0031-9422(80)85121-1

30. Du, J., Meng, L., Qiu, M., Chen, S., Zhang, B., Song, W., Cong, P., & Zheng, X. (2022). Ammonia-oxidizing archaea and ammonia-oxidizing bacteria communities respond differently in oxy-gen-limited habitats. Frontiers in Environmental Science, 10. https://www.frontiersin.org/articles/10.3389/fenvs.2022.976618

31. Elling, F. J., Könneke, M., Lipp, J. S., Becker, K. W., Gagen, E. J., & Hinrichs, K.-U. (2014). Effects of growth phase on the membrane lipid composition of the thaumarchaeon *Nitrosopumilus maritimus* and their implications for archaeal lipid distributions in the marine environment. Geochimica et Cosmochimica Acta, 141, 579–597. 10.1016/j.gca.2014.07.005

32. Elling, F. J., Könneke, M., Mußmann, M., Greve, A., & Hinrichs, K.-U. (2015). Influence of temperature, pH, and salinity on membrane lipid composition and TEX86 of marine planktonic thaumarchaeal isolates. Geochimica et Cosmochimica Acta, 171, 238–255. 10.1016/j.gca.2015.09.004

33. Elling, F. J., Könneke, M., Nicol, G. W., Stieglmeier, M., Bayer, B., Spieck, E., de la Torre, J. R., Becker, K. W., Thomm, M., Prosser, J. I., Herndl, G. J., Schleper, C., & Hinrichs, K.-U. (2017). Chemotaxonomic characterisation of the thaumarchaeal lipidome. Environmental Microbiology, 19(7), 2681–2700. 10.1111/1462-2920.13759

34. Evans, T. W., Könneke, M., Lipp, J. S., Adhikari, R. R., Taubner, H., Elvert, M., & Hinrichs, K.-U. (2018). Lipid biosynthesis of Nitrosopumilus maritimus dissected by lipid specific radioisotope probing (lipid-RIP) under contrasting ammonium supply. Geochimica et Cosmochimica Acta, 242, 51–63. 10.1016/j.gca.2018.09.001

35. Feyhl-Buska, J., Chen, Y., Jia, C., Wang, J.-X., Zhang, C. L., & Boyd, E. S. (2016). Influence of Growth Phase, pH, and Temperature on the Abundance and Composition of Tetraether Lipids in the Thermoacidophile Picrophilus torridus. Frontiers in Microbiology, 7. 10.3389/fmicb.2016.01323

36. Fournier, R. O., Weltman, U., Counce, D., White, L. D., & Janik, C. J. (2002). Results of weekly chemical and isotopic monitoring of selected springs in Norris Geyser Basin, Yellowstone National Park during June-September, 1995. In Results of weekly chemical and isotopic monitoring of selected springs in Norris Geyser Basin, Yellowstone National Park during June-September, 1995 (USGS Numbered Series No. 2002–344; Open-File Report, Vols. 2002–344, p. 50). U.S. Geological Survey. 10.3133/ofr02344

37. Gabriel, J. L., & Lee Gau Chong, P. (2000). Molecular modeling of archaebacterial bipolar tetraether lipid membranes. Chemistry and Physics of Lipids, 105(2), 193–200. 10.1016/S0009-3084(00)00126-2

38. Grossman, E. L., & Joachimski, M. M. (2022). Ocean temperatures through the Phanerozoic reassessed. Scientific Reports, 12(1), 8938. 10.1038/s41598-022-11493-1

39. Harvey, H. R., Fallon, R. D., & Patton, J. S. (1986). The effect of organic matter and oxygen on the degradation of bacterial membrane lipids in marine sediments. Geochimica et Cosmochimica Acta, 50(5), 795–804. 10.1016/0016-7037(86)90355-8

40. Hatzenpichler, R., Lebedeva, E. V., Spieck, E., Stoecker, K., Richter, A., Daims, H., & Wagner, M. (2008). A moderately thermophilic ammonia-oxidizing crenarchaeote from a thermal spring. Proceedings of the National Academy of Sciences, 105(6), 2134–2139. 10.1073/pnas.0708857105

41. He, L., Zhang, C. L., Dong, H., Fang, B., & Wang, G. (2012). Distribution of glycerol dialkyl glycerol tetraethers in Tibetan thermal springs. Geoscience Frontiers, 3(3), 289–300. 10.1016/j.gsf.2011.11.015

42. Heasler, H., & Jaworowski, C. (2018). Hydrothermal monitoring of Norris Geyser Basin, Yellowstone National Park, USA, using airborne thermal infrared imagery. Geothermics, 72, 24–46. 10.1016/j.geothermics.2017.10.016

43. Heeter, K. J., Rochner, M. L., & Harley, G. L. (2021). Summer Air Temperature for the Greater Yellowstone Ecoregion (770–2019 CE) Over 1,250 Years. Geophysical Research Letters, 48(7), e2020GL092269. 10.1029/2020GL092269

44. Hou, W., Wang, S., Dong, H., Jiang, H., Briggs, B. R., Peacock, J. P., Huang, Q., Huang, L., Wu, G., Zhi, X., Li, W., Dodsworth, J. A., Hedlund, B. P., Zhang, C., Hartnett, H. E., Dijkstra, P., & Hungate, B. A. (2013). A Comprehensive Census of Microbial Diversity in Hot Springs of Tengchong, Yunnan Province China Using 16S rRNA Gene Pyrosequencing. PLOS ONE, 8(1), e53350. 10.1371/journal.pone.0053350

45. Huguet, C., Hopmans, E. C., Febo-Ayala, W., Thompson, D. H., Sinninghe Damsté, J. S., & Schouten, S. (2006). An improved method to determine the absolute abundance of glycerol dibiphytanyl glycerol tetraether lipids. Organic Geochemistry, 37(9), 1036–1041. 10.1016/j.orggeochem.2006.05.008

46. Hurley, S. J., Elling, F. J., Könneke, M., Buchwald, C., Wankel, S. D., Santoro, A. E., Lipp, J. S., Hinrichs, K.-U., & Pearson, A. (2016). Influence of ammonia oxidation rate on thaumarchaeal lipid composition and the TEX86 temperature proxy. Proceedings of the National Academy of Sciences, 113(28), 7762–7767. 10.1073/pnas.1518534113

47. Hurley, S. J., Lipp, J. S., Close, H. G., Hinrichs, K.-U., & Pearson, A. (2018). Distribution and export of isoprenoid tetraether lipids in suspended particulate matter from the water column of the Western Atlantic Ocean. Organic Geochemistry, 116, 90–102. 10.1016/j.orggeochem.2017.11.010

48. Hurwitz, S., & Lowenstern, J. B. (2014). Dynamics of the Yellowstone hydrothermal system. Reviews of Geophysics, 52(3), 375–411. 10.1002/2014RG000452

49. James, C. N., Copeland, R., and Lytle, D.A.* (2004). Relationships between oxidation-reduction potential, oxidant, and pH in drinking water. Presented at AWWA Water Quality Technology Conference, San Antonio, TX.

50. Jia, C., Zhang, C. L., Xie, W., Wang, J.-X., Li, F., Wang, S., Dong, H., Li, W., & Boyd, E. S. (2014). Differential temperature and pH controls on the abundance and composition of H-GDGTs in terrestrial thermal springs. Organic Geochemistry, 75, 109–121. 10.1016/j.orggeochem.2014.06.009

51. Jiang, L.-Q., Carter, B. R., Feely, R. A., Lauvset, S. K., & Olsen, A. (2019). Surface ocean pH and buffer capacity: Past, present and future. Scientific Reports, 9(1), Article 1. 10.1038/s41598-019-55039-4

52. Justnes, H. (2020). Transformation Kinetics of Burnt Lime in Freshwater and Sea Water. Materials, 13. 10.3390/ma13214926

53. Kato, S., Itoh, T., Yuki, M., Nagamori, M., Ohnishi, M., Uematsu, K., Suzuki, K., Takashina, T., & Ohkuma, M. (2019). Isolation and characterization of a thermophilic sulfur– and iron-reducing thaumarchaeote from a terrestrial acidic thermal spring. The ISME Journal, 13(10), 2465–2474. 10.1038/s41396-019-0447-3

54. Kaur, G., Mountain, B. W., Stott, M. B., Hopmans, E. C., & Pancost, R. D. (2015). Temperature and pH control on lipid composition of silica sinters from diverse thermal springs in the Taupo Volcanic Zone, New Zealand. Extremophiles, 19(2), 327–344. 10.1007/s00792-014-0719-9

55. Kim, J.-H., Schouten, S., Hopmans, E. C., Donner, B., & Sinninghe Damsté, J. S. (2008). Global sediment core-top calibration of the TEX86 paleothermometer in the ocean. Geochimica et Cosmochimica Acta, 72(4), 1154–1173. 10.1016/j.gca.2007.12.010

56. Kim, J.-H., van der Meer, J., Schouten, S., Helmke, P., Willmott, V., Sangiorgi, F., Koç, N., Hopmans, E. C., & Damsté, J. S. S. (2010). New indices and calibrations derived from the distribution of crenarchaeal isoprenoid tetraether lipids: Implications for past sea surface temperature reconstructions. Geochimica et Cosmochimica Acta, 74(16), 4639–4654. 10.1016/j.gca.2010.05.027

57. Konings, W. N., Albers, S.-V., Koning, S., & Driessen, A. J. M. (2002). The cell membrane plays a crucial role in survival of bacteria and archaea in extreme environments. Antonie van Leeuwenhoek, 81(1), 61–72. 10.1023/A:1020573408652

58. Könneke, M., Bernhard, A. E., de la Torre, J. R., Walker, C. B., Waterbury, J. B., & Stahl, D. A. (2005). Isolation of an autotrophic ammonia-oxidizing marine archaeon. Nature, 437(7058), 543–546. 10.1038/nature03911

59. Kuypers, M. M. M., Blokker, P., Erbacher, J., Kinkel, H., Pancost, R. D., Schouten, S., & Sinninghe Damsté, J. S. (2001). Massive Expansion of Marine Archaea During a Mid-Cretaceous Oceanic Anoxic Event. Science, 293(5527), 92–95. 10.1126/science.1058424

60. Li, F., Zhang, C. L., Dong, H., Li, W., & Williams, A. (2013). Environmental controls on the distribution of archaeal lipids in Tibetan thermal springs: Insight into the application of organic proxies for biogeochemical processes. Environmental Microbiology Reports, 5(6), 868–882. 10.1111/1758-2229.12089

61. Lipp, J. S., & Hinrichs, K.-U. (2009). Structural diversity and fate of intact polar lipids in marine sediments. Geochimica et Cosmochimica Acta, 73(22), 6816–6833. 10.1016/j.gca.2009.08.003

62. Liu, X., Lipp, J. S., & Hinrichs, K.-U. (2011). Distribution of intact and core GDGTs in marine sediments. Organic Geochemistry, 42(4), 368–375. 10.1016/j.orggeochem.2011.02.003

63. Lloyd, C. T., Iwig, D. F., Wang, B., Cossu, M., Metcalf, W. W., Boal, A. K., & Booker, S. J. (2022). Discovery, structure and mechanism of a tetraether lipid synthase. Nature, 609(7925), Article 7925. 10.1038/s41586-022-05120-2

64. Luo, Z.-H., Li, Q., Xie, Y.-G., Lv, A.-P., Qi, Y.-L., Li, M.-M., Qu, Y.-N., Liu, Z.-T., Li, Y.-X., Rao, Y.-Z., Jiao, J.-Y., Liu, L., Narsing Rao, M. P., Hedlund, B. P., Evans, P. N., Fang, Y., Shu, W.-S., Huang, L.-N., Li, W.-J., & Hua, Z.-S. (2024). Temperature, pH, and oxygen availability contributed to the functional differentiation of ancient Nitrososphaeria. The ISME Journal, 18(1), wrad031. 10.1093/ismejo/wrad031

65. Macalady, J. L., Vestling, M. M., Baumler, D., Boekelheide, N., Kaspar, C. W., & Banfield, J. F. (2004). Tetraether-linked membrane monolayers in Ferroplasma spp: A key to survival in acid. Extremophiles, 8(5), 411–419. 10.1007/s00792-004-0404-5

66. National Institute of Standard and Technology. 4.1.4.4. LOESS (aka LOWESS) – Introduction to Process Modeling – Engineering Statistics Handbook. NIST.gov. https://www.itl.nist.gov/div898/handbook/pmd/section1/pmd144.htm

67. Neale, C. M. U., Jaworowski, C., Heasler, H., Sivarajan, S., & Masih, A. (2016). Hydrothermal monitoring in Yellowstone National Park using airborne thermal infrared remote sensing. Remote Sensing of Environment, 184, 628–644. 10.1016/j.rse.2016.04.016

68. Oger, P. M., & Cario, A. (2013). Adaptation of the membrane in Archaea. Biophysical Chemistry, 183, 42–56. 10.1016/j.bpc.2013.06.020

69. Palmer, M. R., Pearson, P. N., & Cobb, S. J. (1998). Reconstructing Past Ocean pH-Depth Profiles. Science, 282(5393), 1468–1471. 10.1126/science.282.5393.1468

70. Pancost, R. D., Pressley, S., Coleman, J. M., Talbot, H. M., Kelly, S. P., Farrimond, P., Schouten, S., Benning, L., & Mountain, B. W. (2006). Composition and implications of diverse lipids in New Zealand Geothermal sinters. Geobiology, 4(2), 71–92. 10.1111/j.1472-4669.2006.00069.x

71. Paraiso, J. J., Williams, A. J., Huang, Q., Wei, Y., Dijkstra, P., Hungate, B., Dong, H., Hedlund, B., & Zhang, C. (2013). The distribution and abundance of archaeal tetraether lipids in U.S. Great Basin thermal springs. Frontiers in Microbiology, 4. https://www.frontiersin.org/articles/10.3389/fmicb.2013.00247

72. Payne, D., Dunham, E. C., Mohr, E., Miller, I., Arnold, A., Erickson, R., Fones, E. M., Lindsay, M. R., Colman, D. R., & Boyd, E. S. (2019). Geologic legacy spanning >90 years explains unique Yellowstone hot spring geochemistry and biodiversity. Environmental Microbiology, 21(11), 4180– 4195. 10.1111/1462-2920.14775

73. Pearson, A., Huang, Z., Ingalls, A. E., Romanek, C. S., Wiegel, J., Freeman, K. H., Smittenberg, R. H., & Zhang, C. L. (2004). Nonmarine Crenarchaeol in Nevada thermal Springs. Applied and Environmental Microbiology, 70(9), 5229–5237. 10.1128/AEM.70.9.5229-5237.2004

74. Pearson, A., Pi, Y., Zhao, W., Li, W., Li, Y., Inskeep, W., Perevalova, A., Romanek, C., Li, S., & Zhang, C. L. (2008). Factors Controlling the Distribution of Archaeal Tetraethers in Terrestrial thermal Springs. Applied and Environmental Microbiology, 74(11), 3523–3532. 10.1128/AEM.02450-07

75. Pearson, A., & Ingalls, A. E. (2013). Assessing the Use of Archaeal Lipids as Marine Environmental Proxies. Annual Review of Earth and Planetary Sciences, 41(1), 359–384. 10.1146/annurev-earth-050212-123947

76. Pitcher, A., Hopmans, E. C., Mosier, A. C., Park, S.-J., Rhee, S.-K., Francis, C. A., Schouten, S., & Sinninghe Damsté, J. S. (2011). Core and Intact Polar Glycerol Dibiphytanyl Glycerol Tetraether Lipids of Ammonia-Oxidizing Archaea Enriched from Marine and Estuarine Sediments. Applied and Environmental Microbiology, 77(10), 3468–3477. 10.1128/AEM.02758-10

77. Pitcher, A., Rychlik, N., Hopmans, E. C., Spieck, E., Rijpstra, W. I. C., Ossebaar, J., Schouten, S., Wagner, M., & Sinninghe Damsté, J. S. (2010). Crenarchaeol dominates the membrane lipids of Candidatus Nitrososphaera gargensis, a thermophilic Group I.1b Archaeon. The ISME Journal, 4(4), 542–552. 10.1038/ismej.2009.138

78. Pitcher, A., Schouten, S., & Sinninghe Damsté, J. S. (2009). In Situ Production of Crenarchaeol in Two California thermal Springs. Applied and Environmental Microbiology, 75(13), 4443–4451. 10.1128/AEM.02591-08

79. Podar, P. T., Yang, Z., Björnsdóttir, S. H., & Podar, M. (2020). Comparative Analysis of Microbial Diversity Across Temperature Gradients in Hot Springs From Yellowstone and Iceland. Frontiers in Microbiology, 11. 10.3389/fmicb.2020.01625

80. Qin, W., Carlson, L. T., Armbrust, E. V., Devol, A. H., Moffett, J. W., Stahl, D. A., & Ingalls, A. E. (2015). Confounding effects of oxygen and temperature on the TEX86 signature of marine Thaumarchaeota. Proceedings of the National Academy of Sciences, 112(35), 10979–10984. 10.1073/pnas.1501568112

81. R Core Team (2024). R: A language and environment for statistical computing. R Foundation for Statistical Computing. https://www.r-project.org/

82. Robert, F., & Chaussidon, M. (2006). A palaeotemperature curve for the Precambrian oceans based on silicon isotopes in cherts. Nature, 443(7114), Article 7114. 10.1038/nature05239

83. Schouten, S., Hopmans, E. C., Pancost, R. D., & Damsté, J. S. S. (2000). Widespread occurrence of structurally diverse tetraether membrane lipids: Evidence for the ubiquitous presence of low-temperature relatives of hyperthermophiles. Proceedings of the National Academy of Sciences, 97(26), 14421–14426. 10.1073/pnas.97.26.14421

84. Schouten, S., Hopmans, E. C., Schefuß, E., & Sinninghe Damsté, J. S. (2002). Distributional variations in marine crenarchaeotal membrane lipids: A new tool for reconstructing ancient sea water temperatures? Earth and Planetary Science Letters, 204(1), 265–274. 10.1016/S0012-821X(02)00979-2

85. Schouten, S., Hopmans, E. C., & Sinninghe Damsté, J. S. (2013). The organic geochemistry of glycerol dialkyl glycerol tetraether lipids: A review. Organic Geochemistry, 54, 19–61. 10.1016/j.orggeochem.2012.09.006

86. Schouten, S., van der Meer, M. T. J., Hopmans, E. C., Rijpstra, W. I. C., Reysenbach, A.-L., Ward, D. M., & Sinninghe Damsté, J. S. (2007). Archaeal and Bacterial Glycerol Dialkyl Glycerol Tetraether Lipids in thermal Springs of Yellowstone National Park. Applied and Environmental Microbiology, 73(19), 6181–6191. 10.1128/AEM.00630-07

87. Shock, E. L., Holland, M., Meyer-Dombard, D., Amend, J. P., Osburn, G. R., & Fischer, T. P. (2010). Quantifying inorganic sources of geochemical energy in hydrothermal ecosystems, Yellowstone National Park, USA. Geochimica et Cosmochimica Acta, 74(14), 4005–4043. 10.1016/j.gca.2009.08.036

88. Sinninghe Damsté, J. S., Rijpstra, W. I. C., Hopmans, E. C., den Uijl, M. J., Weijers, J. W. H., & Schouten, S. (2018). The enigmatic structure of the crenarchaeol isomer. Organic Geochemistry, 124, 22–28. 10.1016/j.orggeochem.2018.06.005

89. Sinninghe Damsté, J. S., Rijpstra, W. I. C., Hopmans, E. C., Jung, M.-Y., Kim, J.-G., Rhee, S.-K., Stieglmeier, M., & Schleper, C. (2012). Intact Polar and Core Glycerol Dibiphytanyl Glycerol Tetraether Lipids of Group I.1a and I.1b Thaumarchaeota in Soil. Applied and Environmental Microbiology, 78(19), 6866–6874. 10.1128/AEM.01681-12

90. Sinninghe Damsté, J. S., Schouten, S., Hopmans, E. C., Duin, A. C. T. van, & Geenevasen, J. A. J. (2002). Crenarchaeol. Journal of Lipid Research, 43(10), 1641–1651. 10.1194/jlr.M200148-JLR200

91. Slonczewski, J. L., Fujisawa, M., Dopson, M., & Krulwich, T. A. (2009). Cytoplasmic pH Measurement and Homeostasis in Bacteria and Archaea. In R. K. Poole (Ed.), Advances in Microbial Physiology (Vol. 55, pp. 1–317). Academic Press. 10.1016/S0065-2911(09)05501-5

92. Stahl, D. A., & de la Torre, J. R. (2012). Physiology and Diversity of Ammonia-Oxidizing Archaea. Annual Review of Microbiology, 66(1), 83–101. 10.1146/annurev-micro-092611-150128

93. Taylor, K. W. R., Huber, M., Hollis, C. J., Hernandez-Sanchez, M. T., & Pancost, R. D. (2013). Re-evaluating modern and Palaeogene GDGT distributions: Implications for SST reconstructions. Global and Planetary Change, 108, 158–174. 10.1016/j.gloplacha.2013.06.011

94. Tierney, J. E. (2012). GDGT Thermometry: Lipid Tools for Reconstructing Paleotemperatures. The Paleontological Society Papers, 18, 115–132. 10.1017/S1089332600002588

95. Tourte, M., Schaeffer, P., Grossi, V., & Oger, P. M. (2022). Membrane adaptation in the hyperthermophilic archaeon Pyrococcus furiosus relies upon a novel strategy involving glycerol monoalkyl glycerol tetraether lipids. Environmental Microbiology, 24(4), 2029–2046. 10.1111/1462-2920.15923

96. Valentine, D. L. (2007). Adaptations to energy stress dictate the ecology and evolution of the Archaea. Nature Reviews Microbiology, 5(4), Article 4. 10.1038/nrmicro1619

97. van de Vossenberg, J. L., Driessen, A. J., & Konings, W. N. (1998). The essence of being extremophilic: The role of the unique archaeal membrane lipids. Extremophiles: Life Under Extreme Conditions, 2(3), 163–170. 10.1007/s007920050056

98. Weber, Y., Sinninghe Damsté, J. S., Hopmans, E. C., Lehmann, M. F., & Niemann, H. (2017). Incomplete recovery of intact polar glycerol dialkyl glycerol tetraethers from lacustrine suspended biomass. Limnology and Oceanography: Methods, 15(9), 782–793. 10.1002/lom3.10198

99. White, D. C., Davis, W. M., Nickels, J. S., King, J. D., & Bobbie, R. J. (1979). Determination of the Sedimentary Microbial Biomass by Extractible Lipid Phosphate. Oecologia, 40(1), 51–62.

100. Wu, W., Zhang, C., Wang, H., He, L., Li, W., & Dong, H. (2013). Impacts of temperature and pH on the distribution of archaeal lipids in Yunnan thermal springs, China. Frontiers in Microbiology, 4. https://www.frontiersin.org/articles/10.3389/fmicb.2013.00312

101. Xie, W., Zhang, C. L., Wang, J., Chen, Y., Zhu, Y., de la Torre, J. R., Dong, H., Hartnett, H. E., Hedlund, B. P., & Klotz, M. G. (2015). Distribution of ether lipids and composition of the archaeal community in terrestrial geothermal springs: Impact of environmental variables. Environmental Microbiology, 17(5), 1600–1614. 10.1111/1462-2920.12595

102. Yang, Y., Zhang, C., Lenton, T. M., Yan, X., Zhu, M., Zhou, M., Tao, J., Phelps, T. J., & Cao, Z. (2021). The Evolution Pathway of Ammonia-Oxidizing Archaea Shaped by Major Geological Events. Molecular Biology and Evolution, 38(9), 3637–3648. 10.1093/molbev/msab129

103. Zeng, Z., Chen, H., Yang, H., Chen, Y., Yang, W., Feng, X., Pei, H., & Welander, P. V. (2022). Identification of a protein responsible for the synthesis of archaeal membrane-spanning GDGT lipids. Nature Communications, 13(1), Article 1. 10.1038/s41467-022-29264-x

104. Zhang, C. L., Pearson, A., Li, Y.-L., Mills, G., & Wiegel, J. (2006). Thermophilic Temperature Optimum for Crenarchaeol Synthesis and Its Implication for Archaeal Evolution. Applied and Environmental Microbiology, 72(6), 4419–4422. 10.1128/AEM.00191-06

105. Zhang, Y. G., Pagani, M., & Wang, Z. (2016). Ring Index: A new strategy to evaluate the integrity of TEX86 paleothermometry. Paleoceanography, 31(2), 220–232. 10.1002/2015PA002848

106. Zhao, W., Song, Z., Jiang, H., Li, W., Mou, X., Romanek, C. S., Wiegel, J., Dong, H., & Zhang, C. L. (2011). Ammonia-oxidizing Archaea in Kamchatka thermal Springs. Geomicrobiology Journal, 28(2), 149–159. 10.1080/01490451003753076

107. Zhou, A., Weber, Y., Chiu, B. K., Elling, F. J., Cobban, A. B., Pearson, A., & Leavitt, W. D. (2020). Energy flux controls tetraether lipid cyclization in Sulfolobus acidocaldarius. Environmental Microbiology, 22(1), 343–353. 10.1111/1462-2920.14851

